# Parthenogenesis, sexual conflict, and selection on fertilization rates in switching environments

**DOI:** 10.1101/2023.06.09.544312

**Authors:** Xiaoyuan Liu, Jon W. Pitchford, George W.A. Constable

## Abstract

In the face of varying environments, organisms exhibit a variety of reproductive modes, from asexuality to obligate sexuality. Should reproduction be sexual, the morphology of the sex cells (gametes) produced by these organisms has important evolutionary implications; these cells can be the same size (isogamy), one larger and one smaller (anisogamy), and finally the larger cell can lose its capacity for motility (oogamy, the familiar sperm-egg system). Understanding the origin of the sexes, which lies in the types of gametes they produce, thus amounts to explaining these evolutionary transitions. Here we extend classic results in this area by exploring these transitions in a model in which organisms can reproduce both sexually and asexually (a reproductive mode present in many algae). In particular, we investigate the co-evolution of gamete cell size with fertilization rate, which is a proxy for motility and pheromone production but usually held constant in such models. Using adaptive dynamics generalized to the case of switching environments, we find that isogamy can evolve to anisogamy through evolutionary branching, and that anisogamy can evolve to oogamy or suppressed pheromone production through a further branching driven by sexual conflict. We also derive analytic conditions on the model parameters required to arrest evolution on this isogamy-oogamy trajectory, with low fertilization rates and stochastically switching environments stabilizing isogamy under a bet-hedging strategy, and low fertilization costs stabilizing anisogamy and pheromone production.

## 1. Introduction

The origin of the sexes lies in the types of sex cells they produce; males produce small microgametes while females produce larger macrogametes [1]. This gamete size dimorphism is referred to as a state of anisogamy [2]. Should the microgamete be motile (sperm) and the macrogamete sessile (egg), the population is said to be oogamous [3]. While anisogamy is the most commonly observed mode of sexual reproduction in eukaryotic organisms [4], having evolved several times in evolutionary history [5], it is now widely accepted to be derived from isogamy (equal gamete sizes) [6]. While rarer, isogamous species include study organisms such as the yeast *Saccharomyces cerevisiae* [7] and the green algae *Chlamydomonas reinhardtii* [8], where self-incompatible mating types play the role of ancestral versions of the true sexes [9]. Indeed, the volvocine algae (of which *C. reinhardtii* is a member), provide neat empirical examples of these transitions, with phylogenetic analysis indicating that numerous independent lineages have undergone the transition from isogamy, to anisogamy, and finally to oogamy [3, 10]. Explaining the evolutionary mechanisms behind these transitions has been the focus of much work in evolutionary theory.

While theoretical investigations of the evolution of anisogamy date back to the 1930’s [11] and were developed into the 1960’s [12, 13], it was arguably the Parker-Baker-Smith (PBS) model [14] that synthesised these ideas into a complete evolutionary model that is now widely accepted as providing an explanation for the evolution of anisogamy [15, 16, 17]. They assumed that a fixed mass or energy budget is allocated by individuals to gamete production, such that microgametes can be produced in larger quantities than macrogametes. However, while microgametes may be more numerous, they contribute a lower fitness than macrogametes to a fertilized zygote due to their low cytoplasmic volume [18]. In this way the PBS model was able to show how anisogamy was the result of a quality-quantity trade-off, with disruptive selection acting on gamete size [15].

The key elements of the PBS model have since been set in a game theoretic [19] and population genetic [20, 21] context, as well as extended to account for more general reproductive modes such as hermaphrodism [22]. Models using adaptive dynamics [23] in particular have been useful. It has been shown analytically that even in the absence of mating types, anisogamy can evolve from isogamy through evolutionary branching in mass [24] and that this is a stable state. Meanwhile, accounting for self-incompatibility of mating types and investigating the effect of varying fertilization rate, [25] showed that anisogamy can evolve from isogamy through both gamete competition and gamete limitation. Altogether, the results described above suggest that fertilization rate is a crucial factor that may impact gamete survival, and selection is likely to act on the fertilization rate between microgametes and macrogametes.

As addressed, oogamy, the loss of motility in eggs and specialization for motility in sperm, is often seen as the “last step in the evolution of the egg–sperm dichotomy” [26], a view supported by empirical analyses that suggest oogamy is derived from anisogamy [27]. However the theory of this transition is comparatively less studied than the earlier transition from isogamy to anisogamy. Ghiselin [28] provided an argument based on the physiological division of labour between macro and microgametes, with females specialising in provisioning and males in motility. Most other work has considered the evolution of oogamy as a strategy to maximise gamete encounter rate. This can be achieved by having a population of pheromone emitters and receivers [29, 30] and by having a large stationary target egg and small motile sperm [31, 32]. Although the assumption of an inverse speed-size relationship in [32] has justification in some gametic systems, it is also worth noting that positive speed-size relationship have been observed in *C. reinhardtii*, due to larger cells having greater propulsive forces; at scales such as these the precise speed-size relationship is complicated by details of cell morphology [33]. Important theoretical progress was therefore made in [34], which investigated how investment in motility can differ between males and females under differing levels of gamete limitation and different speed-size relationships. Under both positive and negative speed-size relationships, the level of motility investment is biased towards one sex. Strikingly, increasing gamete limitation can trigger a switch from the classic male biased motility investment (oogamy) to female biased motility investment. Lastly, the prior evolution of internal fertilization has been proposed as a mechanism that could generate selection for oogamy [35], consistent with empirical evidence from *volvox* (external fertilization and anisogamous) and its sister lineage *platydorina* (internal fertilization and oogamous) [36].

The work described above all assumes obligate sexual reproduction (unfertilized gametes die at the end of each generation) and a static environment [19, 25, 26]. Recently, however, inspired by the life-histories of green and brown algae such as *Blidingia minima* (isogamous [37]), *Urospora neglecta* (anisogamous [38]), and *Saccharina japonica* (oogamous [39]), whose gametes can develop asexually through parthenogenesis should they fail to find a mate (see [37, 38, 40], respectively), the first of these two assumptions was relaxed in two theoretical papers [41, 42]. In [41], extra survival costs were incurred by gametes developing parthenogenetically, while in [42] extra survival costs on either the parthenogenetic or the sexual reproductive route were considered. They found that in the presence of two self-incompatible mating types, isogamy can be stabilized under low costs to parthenogenesis, while anisogamy is the evolutionarily stable state when fertilization is favoured.

The studies described above on parthenogenesis and the evolution of anisogamy lead to a natural question; if the sexual route (via fertilization) and asexual route (via parthenogenesis) carry different survival costs, how should the fertilization rate evolve to account for this? Should this rate increase (to minimise the number of unfertilized gametes taking a potentially perilous route to survival) or should it decrease (to avoid potential costs incurred during cell fusion)? The modelling of fertilization kinetics is an interesting topic in its own right [43, 44], and plays a role in models for the evolution of anisogamy from isogamy [25]. However fertilization kinetics are also clearly a key element of the evolution of oogamy, with selection on sperm to increase their encounter rate with eggs and selection on eggs to remain sessile.

In this paper, we modify the adaptive dynamics models of [24, 41, 42] to study the co-evolution of gamete size and fertilization rate in species capable of parthenogenesis under external fertilization. For simplicity, we assume an absence of self-incompatible mating types [6, 24]. We show that under such assumptions, anisogamy can evolve from an initial state of isogamy, followed by the subsequent evolution to oogamy under sexual conflict between microgametes and macrogametes. In Section 2, we introduce the model, describing its key behaviour in Section 3. These behaviours include the evolution of oogamy (Section 3.2), conditions that stabilize isogamy (Section 3.3), conditions that stabilize anisogamy (Section 3.4), and the emergence of isogamy as a bet-hedging strategy in switching environments (Section 3.5). Finally in Section 4 we conclude by discussing the results and how they may relate to empirical examples of geographic parthenogenesis [45, 46] in aquatic external fertilizers.

## 2. Model

In this section we describe the specifics of the models we use, paying careful attention to the various time scales involved. We begin by considering the evolutionary model in a fixed environment (Section 2.1), before generalizing to the case of a under which bet-hedging strategies can evolve (Section 2.2).

### 2.1. Model dynamics in a fixed environment

The evolutionary dynamics of the model are built from a hierarchy of timescales, which are particularly important to keep in mind once environmental switching is introduced in later sections. The shortest timescale is the generational timescale. The intermediate timescale is that over which the invasion of a rare mutant (taking place over many generations) can occur. The longest timescale is the evolutionary timescale, representing the cumulative effect of multiple mutations and invasions.

#### Dynamics within each generation

At the start of each discrete generation, a number of adults with haploid mass (or energy budget) *M* produce gametes (mass *m*), such that mass/energy budget is conserved (i.e. each adult produces *M*/*m* gametes). Note that this implicitly assumes, for simplicity, that a continuous (rather than discrete) number of gametes is possible. Gametes then enter a pool and the fertilization process takes place. After a finite time window, the resultant cells face a round of survival dependent on their mass. The surviving cells form the basis of the next generation, completing the generational cycle, as illustrated in Figure 1.

**Figure 1:**
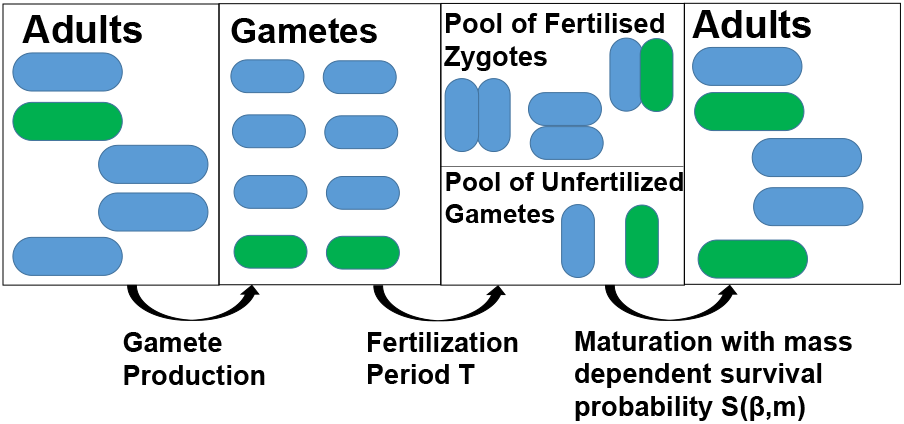
Schematic of dynamics within each generation. Mature cells (adults) produce gametes at the start of a generation. All the gametes are given a fixed time period *T* in which to complete the fertilization process. At the end of the fertilization period, there will be a pool of fertilized zygotes and unfertilized gametes, both of which are capable of maturation. Each cell survives according to its independent survival functions ((1 − *C*_*z*_)*S* (*β, m*_1_ + *m*_2_) and (1 − *C*_*p*_)*S* (*β, m*_*i*_) respectively) to produce a number of mature cells in the subsequent generation. The pool of gametes consist of resident (blue) and mutant gametes (green), where the mutation occurs in either the mass *m* or fertilization rate *α*.

#### Fertilization Kinetics

For simplicity, we assume that all gametes may fertilize each other (i.e. there are no self-incompatible mating types). Given a total of *A* adults, the population is initially comprised of *N* = (*AM*)/*m* single gametes produced by adults through meiosis. We assume fertilization is external, with cells fertilizing according to mass action dynamics at rate *α*, such that the number of single (unfertilized) cells, *N*, is given by the solution to

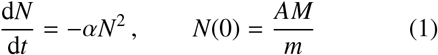

with *α* ≥ 0 (see also [43]). At the end of the fertilization window, which is assumed to have a duration *T*, there are therefore *N*(*T*) single cells remaining, and (*N*(0) − *N*(*T*))/2 fertilized cells.

The parameter *α* is variously referred to as the fertilization rate, the collision rate [43], the “‘aptitude’ for union” [13], or the “bimolecular reaction constant” [47]. We will refer to *α* as the fertilization rate, and treat it as a trait subject to evolution. Overall, *α* captures the compound effect of the propensity for fertilization between cells encountering each other, as well as additional mechanisms to enhance cell encounter rate and fertilization affinity such as increased emission or response to chemoattractants [30], pheromones [48] and gamete recognition proteins [49], or cell motility [50]. In practice there is a likely upper-bound on *α*, brought about by energy trade-offs, diffusion in aquatic environments, or dispersal in terrestrial environments. To account for this we introduce a ceiling on *α*, and restrict its evolutionary dynamics to the range *α*_max_ ≥ *α* ≥ 0.

#### Survival Probability

At the end of a finite fertilization window at time *T*, the population will consist of both fertilized and unfertilized gametes. Fertilized gametes produce diploid zygotes, while unfertilized gametes can develop as haploids parthenogenetically (e.g. parthenosporophytes [51]). We assume that the probability that either of these cell types survives is given by the Vance survival function [18], which is a common assumption in the literature [19, 25, 56]. Given a cell size *m*_*c*_ (for either fertilized or unfertilized cells), the survival probability is given by

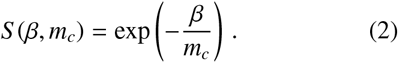

Note that this is an increasing function of cell size, and we do not account for gamete mortality during the fertilization period. Thus, although both fertilized and unfertilized gametes are exposed to the same survival function, fertilized cells (with a mass around twice the size of unfertilized cells in a monomorphic isogamous population) will have a greater survival probability than unfertilized cells. Meanwhile for a given mass *m*_*c*_, increasing *β* will decrease the survival probability. We therefore refer to *β* as the resistance to survival, with high *β* corresponding to harsh environments in which survival is difficult, and low *β* corresponding to more benign environments in which even cells of modest mass have a high probability of surviving.

In addition to the benefits explained in the introduction, fertilization may also carry risks: generally these include cell-fusion failure [57], selfish extra-genomic elements in the cytoplasm [58] and cytoplasmic conflict [59, 60, 61]. However costs may also arise in multicellular organisms if the reproductive output of fertilized diploids (e.g. sporophytes) is less than twice that of unfertilized haploids (e.g. parthenosporophytes). In fact, if the reproductive output of fertilized diploids is equal to that of unfertilized haploids, one obtains a version of the classic twofold cost of sex [52, 53] for species capable of haploid development, as illustrated in Figure 2. In addition, sexual reproduction is time and energy intensive, and requires investing in finding a mate [62]. In light of these varied costs, we introduce an additional, simplified, fixed cost 1 ≥ *C*_*z*_ ≥ 0, applied to fertilized zygotes independent of their mass and fertilization rate.

**Figure 2:**
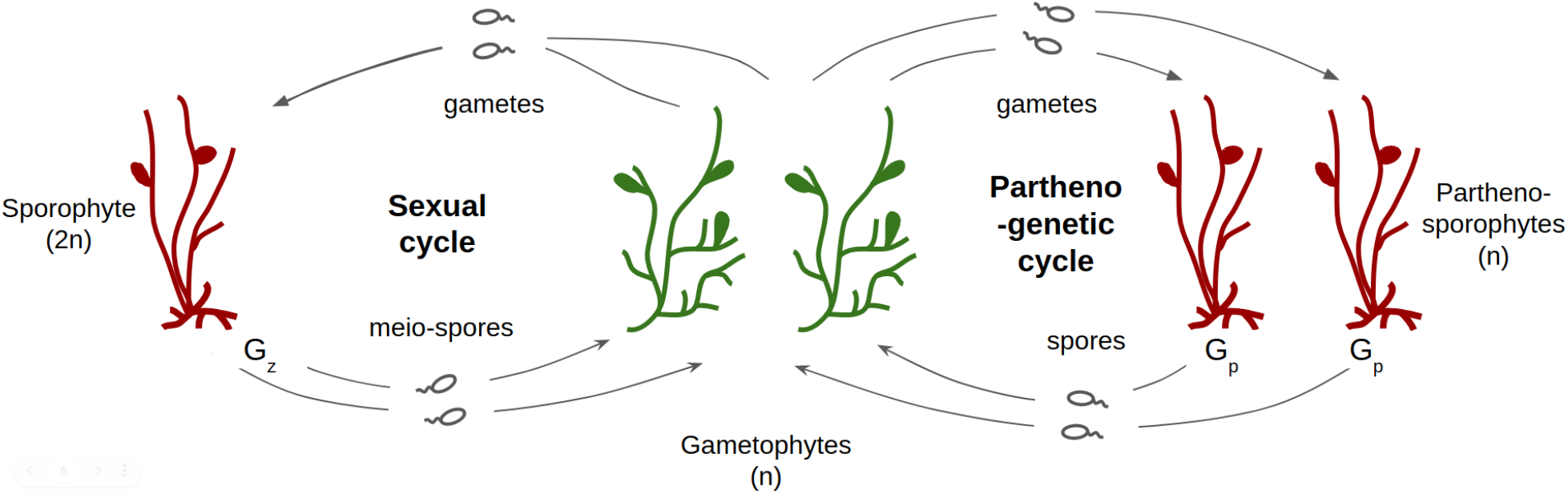
Diagram illustrating some of the potential costs of fertilization, with details inspired by the life-cycle of Ectocarpus [51]. Haploid gametes can become fertilized to form diploid sporophytes. Following genetic recombination, sporophytes produce *G*_*z*_ haploid meio-spores. Assuming Mendelian inheritance, this results in *G*_*z*_/2 meio-spores associated with the genetic traits of each of the gametes contributing to the sporophyte. Conversely haploid gametes can remain unfertilized to become haploid parthenosporophytes. These parthenosporophytes produce *G*_*p*_ haploid spores that inherit the genetic identity of their gametic ancestor. Therefore if *G*_*z*_ = *G*_*p*_, gametes entering the sexual cycle produce half as many meio-spores as gametes entering the parthenogenetic cycle produce spores. This scenario is comparable to the classic twofold cost of sex first described by Williams and Maynard Smith [52, 53] (see also [54, 55]). Should *G*_*z*_ < 2*G*_*p*_ (i.e. sporophytes produce meiospores at anything less than twice the number parthenosporophytes produce spores), a lower, but still present, cost to fertilization persists. Further mathematical details, including the relationship between *G*_*z*_ and *G*_*p*_ and the costs *C*_*z*_ and *C*_*p*_ used in our model, are provided in Appendix A. The figure is adapted from [51] to account for the fact that in our model, both male and female gameteophytes can in principle develop parthenogenetically.

Similarly, there can be costs associated to parthenogenetic development such as reduced fitness due to lack of genetic diversity [63] and the possibility of failure of parthenogenetic development [41]. That parthenogenesis is absent in many algae, while in the green alga green *Monostroma angicava* gametes only reproduce parthenogenetically should they fail to find a mate [64], has motivated the modelling of costs associated with this reproductive pathway [41]. We therefore introduce an additional, again simplified and fixed, cost 1 ≥ *C*_*p*_ ≥ 0, applied to unfertilized gametes independent of their mass and fertilization rate.

The final probability of survival for a zygote formed from the fertilization of two gametes of sizes *m*_1_ and *m*_2_ is then given by (1 − *C*_*z*_) exp [−*β*/(*m*_1_ + *m*_2_)], while the probability of survival for an unfertilized cell of size *m* is (1 − *C*_*p*_) exp [−*β*/*m*].

#### Invasion Dynamics

We assume that haploid gametes are characterised by two genetically determined non-recombining traits; their mass, *m*, and their fertilization rate, *α*. We next consider a monomorphic resident population to which a mutant individual is introduced at rate *µ*. As we will be interested in the effect of changing environments (which may switch multiple times over the course of a mutant invasion) it is necessary for us to construct the invasion dynamics for the mutant. Analytically, constructing these invasion dynamics is only possible when mutations in *m* and *α* occur independently (see Appendix B and Appendix C), and so we consider these cases in the remainder of this section. However in Appendix D we show that the evolutionary dynamics that we obtain are identical to those if we allow mutations both in *m* and *α*.

Suppose the mutant produces gametes of a different mass to its ancestor, *m* ± *δm*, where *δm* represents the size of a mutational step. Under this scenario the mutant may produce more or fewer gametes than its ancestor (see Appendix B.1), but the survival probability of its unfertilized cells (mass *m* ± *δm*), mutant-resident fertilized cells (mass 2*m* ± *δm*), and mutant-mutant fertilized cells (mass 2(*m* ± *δm*)), will also be simultaneously decreased or increased (see Eq. (2) and Appendix B.3). The cumulative effect of this quality-quantity trade-off will either lead to selection for or against the mutant over subsequent generations.

Alternatively, the mutant may engage in an increased or decreased fertilization rate relative to its ancestor, at a rate *α* ± *δα* (see Appendix B.2). Under this scenario the mutant fertilizes with residents at their average fertilization rate (2*α* ± *δα*)/2 and other mutants at a rate *α* ± *δα*. Mutants will either contribute to more or fewer fertilized cells and depending on the resistance to survival, *β*, the costs to fertilization, *C*_*z*_, and the costs to parthenogenesis, *C*_*p*_, may experience a selective advantage over the resident by devoting more of its gametes to one of the reproductive routes (see Appendix B.4).

In order to mathematically characterise the invasion dynamics (which occur over discrete generations), we adopt the classical assumptions of adaptive dynamics (see [65] and Appendix E.1). In particular, we assume that *δm* and *δα* are small, so that we can approximate the dynamics continuously, and successive mutations occur sufficiently rarely that each mutation can equilibrate before a new mutation occurs. Denoting by 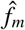 the frequency of mutants of size *m* ± *δm* in the population, and *t*_*g*_ the number of generations, we find (see Appendix C.1)

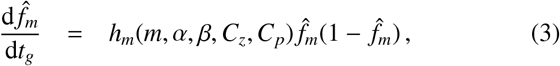

where *h*_*m*_(*m, α, β, C*_*z*_, *C*_*p*_) is a constant selective pressure that depends on the parameters *m, α, β, C*_*z*_, *C*_*p*_ (see Eq. (C.1)). Similarly, denoting by 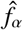 the frequency of mutants with fertilization rate *α* ± *δα* in the population (see Appendix C.2), we find

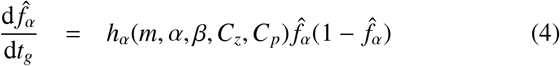

where *h*_*α*_(*m, α, β, C*_*z*_, *C*_*p*_) is a constant that depends on the parameters *m, α, β, C*_*z*_, *C*_*p*_ (see Eq. (C.2)).

We note that although the analysis above shows frequency-independent selection (see Eqs. (3) and (4)), this is only the result of our assumptions that each mutant can equilibriate before any subsequent mutation events [65] and that mutations in *m* and *α* occur independently, which is clearly violated in realistic populations. In particular, the small mutational effects and random occurrence of successive mutational events means that a degree of polymorphism is expected in our model, which can lead to frequency-dependent selection whereby evolutionary branching is a possibility. Indeed frequency-dependent selection does emerge when applying the standard analytical tools of adaptive dynamics [24, 66] that do not require a full calculation of the invasion trajectory (see Appendix D) as well as in our numerical simulations that allow for polymorphic populations (see Appendix E.3).

#### Evolutionary Dynamics

Following standard approaches in adaptive dynamics [67] (see also Appendix E), we construct the evolutionary equations for the gamete mass, *m*, and the fertilization rate, *α*. Denoting by *τ* the evolutionary timescale over which mutations appear and trait substitutions occur, we find that for *α* ≥ 0,

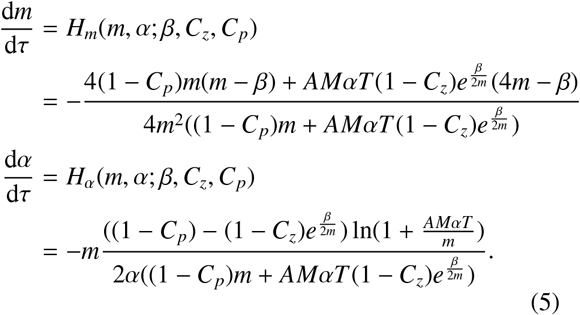

Due to the model setup, *α* can mathematically evolve to be negative from the *α* = 0 state (this would result in an *increase* in gamete numbers during the fertilization period, see Eq. (1)). While mathematically possible, this is clearly not biologically reasonable. We therefore impose *α* = 0 as a biologically realistic (but mathematically discontinuous) boundary. For *α* = 0 and *H*_*α*_(*m*, 0; *β, C*_*z*_, *C*_*p*_) < 0, we then have

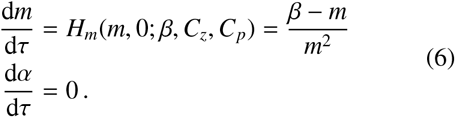

We note that such features are common in simplified models of anisogamy evolution [42], where some evident non-physical behaviour can arise near the boundaries (e.g. when cell masses are zero).

### 2.2. Evolutionary dynamics in switching environments

We now wish to consider the case of a population characterised by gamete mass (*m*) and fertilization rate (*α*) traits, evolving subject to changing environmental conditions. We may again employ techniques from adaptive dynamics, but now generalized to dynamic environments [68]. Explicitly, we allow the resistance to survival to alternate between two values *β*_1_ and *β*_2_; recall from Eq. (2) that if *β*_1_ > *β*_2_ then *β*_1_ represents a comparatively “harsh environment”, where cells have a lower survival probability than in environment *β*_2_.

Switching between these two environments is modelled as a discrete stochastic telegraph process [69, 70]; the time spent in each environment is distributed geometrically (a discrete analogue of the exponential distribution), spending an average period *τ*_1_ ≈ 1/*λ*_1→2_ in environment 1 and *τ*_2_ ≈ 1/*λ*_2→1_ in environment 2, where *λ*_*i*→ *j*_ is the transition rate from environment *i* to *j*. We must carefully consider the magnitude of these timescales in comparison with the other timescales at work in the model (see Section 2.1).

First consider the case where the environmental switching timescales, *τ*_1_ and *τ*_2_, are larger than the generational timescale (*t*_*g*_), but much smaller than the invasion timescale (characterised by the inverse of the strength of selection, proportional to 1/*δm* and 1/*δα*) and the mutational timescale (1/*µ*). We call this the ’fast relative to invasion’ switching regime (FRTI). In this scenario, the population does not switch environments during a single round of fertilization kinetics, but typically switches between the two environmental states many times before an invasion has time to complete.

When switching occurs this frequently, we can approximate the invasion dynamics mathematically by observing that the population experiences the weighted average of the dynamics in the two environments [68] over a large number of generations. Denoting by *P*_1_ and *P*_2_ the probability of finding the population in either of the respective environments, we have

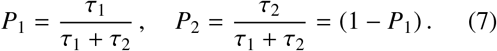

The effective dynamics for the frequency of mutants with mass *m* + *δm* in a resident population of mass *m* during an invasion is then given by

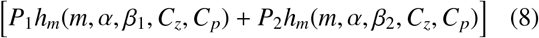

which can be contrasted against the term *h*_*m*_(*m, α, β, C*_*z*_, *C*_*p*_) in Eq. (3). An analogous approach allows us to approximate the invasion dynamics for mutants with a different fertilization rate to their ancestors in this FRTI regime (see Appendix F.1). We note that although these results suggest an algebraic (rather than geometric [71]) averaging of the dynamics across the two environments, the algebraic mean fitness can be obtained as a leading-order approximation of the geometric mean fitness (or long-run growth rate) when variance in the growth rate due to environmental switching is small [72, 73]. We expect this to be the case when switching between environments is fast (*λ*_1→ 2_ and *λ*_2→ 1_ large), as described in Appendix F.6.

With equations for d 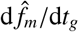 and d 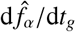 in hand, we can proceed to apply the same standard techniques from adaptive dynamics as were used to derive Eq. (5) from Eqs. (3-4) (see Appendix F). We obtain the effective evolutionary dynamics [68]

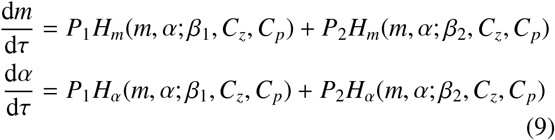

where *H*_*m*_(*m, α*; *β*_1_, *C*_*z*_, *C*_*p*_) and *H*_*α*_(*m, α*; *β*_1_, *C*_*z*_, *C*_*p*_) retain the functional forms given in Eq. (5), and *P*_1_ and *P*_2_ are taken from Eq. (7).

In contrast to the FRTI regime discussed above, we can also investigate the regime in which environmental switching occurs on a comparable or slower timescale than invasion, but still occurs fast relative to the evolutionary timescale. In this scenario, which we call the “fast relative to evolution” switching regime (FRTE), environmental switching occurs on a similar rate to that at which new mutations are introduced, but much faster than the combined effect of mutation and selection (e.g. *λ*_*i*→ *j*_ ≪ *µ* × *δm*), such that only a small number of mutations can fixate in either environment before the population switches to the alternate environment. Although we shall show via simulations in Section 3.5 that this FRTE regime leads to quantitatively different evolutionary trajectories compared to the FRTI regime, we show mathematically in Appendix F.2 that the evolutionary dynamics can be approximated by the same equations (see Eq. (9)).

## 3. Results

In this section we proceed to analyse the evolutionary dynamics derived in Sections 2.1-2.2 and compare our results to numerical simulations of the full stochastic simulations.

### 3.1. Initial evolution of fertilization rate

In Figure 3, we see two potential evolutionary outcomes for the co-evolutionary dynamics of *m* and *α* in a single fixed environment that are dependent on the initial conditions and parameters; the population can either evolve to large fertilization rates (limited only by *α*_max_) or to zero fertilization rates.

**Figure 3:**
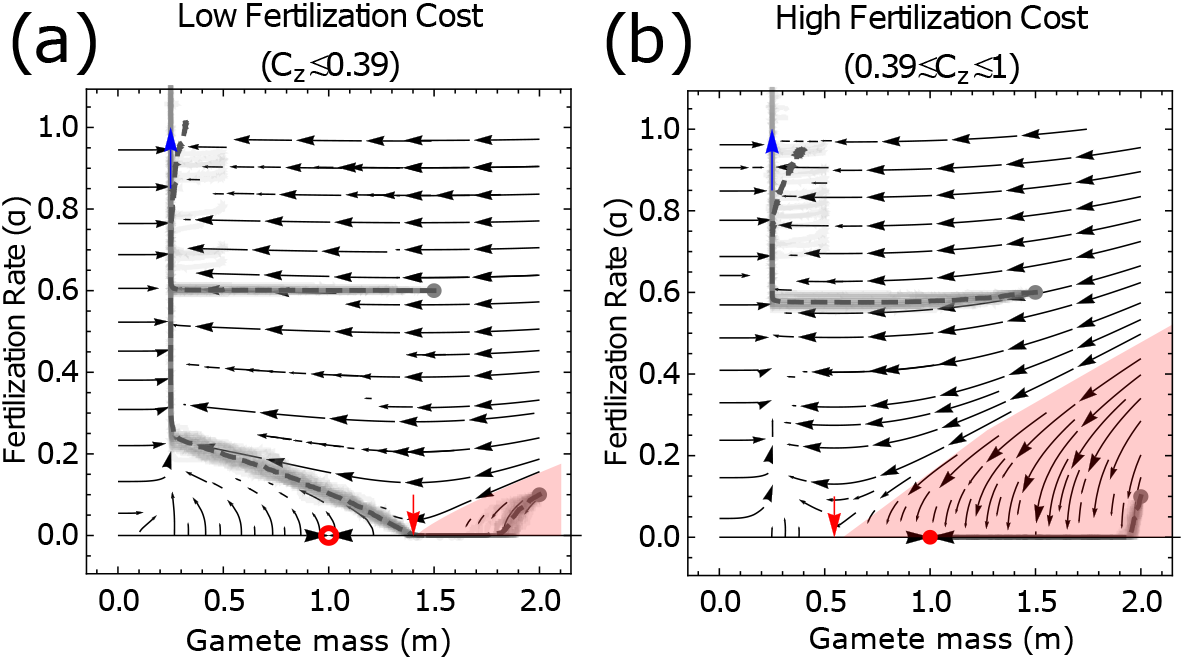
Phase portraits for the co-evolutionary dynamics in a fixed environment (see Eq. (5)). High fertilization rates are the only evolutionary outcome in panel (a), while high and zero fertilization rate are both evolutionary outcomes in panel (b) (see Eq. (10)). The red region shows trajectories leading to points on the α = 0 boundary for which evolution selects for decreasing fertilization rate (dα/dτ < 0) (the point at which dα/dτ = 0 is marked by a red arrow). Red filled (stable) and unfilled (unstable) circles mark a fixed point in the evolutionary dynamics of *m* (*m*^∗^ = *β*, see Eq. (E.2)). Blue arrows illustrate the high fertilization rate fixed point ((*m*^∗^, α^∗^) → (*β*/4, ∞), see Eq. (E.6)). Average population trait trajectories, (⟨*m*⟩ (*t*), ⟨α⟩ (*t*)), from simulation of the stochastic model (see Appendix E.3) are plotted in light gray, and their mean over multiple realisations are given dark gray. The cost to fertilization is *C*_*z*_ = 0.3 (panel (a)) and *C*_*z*_ = 0.6 (panel (b)). In both panels *β* = 1 and *C*_*p*_ = 0. Remaining model parameters are given in Appendix J.

When the cost to fertilization, *C*_*z*_, is low (Figure 3, panel (a)), there exists a smaller region of initial conditions that drive *α* towards zero (red shaded region). When *α* = 0 within this region, selection on gamete mass, *m*, drives the population towards the point *m* = *β* (red dot, see also Appendix E.2). As this point exists outside the region in which d*α*/d*τ* < 0, selection for increased *α* can again manifest along the evolutionary trajectory. Thus when costs are sufficiently low, high fertilization rates are the only evolutionary outcome.

Conversely when costs to fertilization, *C*_*z*_, are intermediate (Figure 3, panel (b)), there exists a larger region of initial conditions that drive *α* towards zero (red shaded region). The point *m* = *β* (red dot), towards which the population evolves when *α* = 0, is now contained within this region in which d*α*/d*τ* < 0, and so is a stable fixed point. Thus when costs are sufficiently high, there are two evolutionary outcomes, depending on the initial conditions; either high fertilization rates or zero fertilization rates.

In Appendix E.2 we conduct a mathematical and numerical analysis to formalise the arguments above. In summary, the possible early evolutionary attractors, (*m*^∗^, *α*^∗^), are given by:

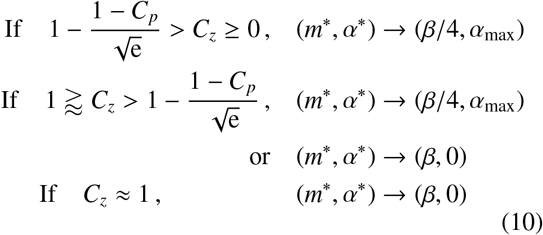

where we note that in the absence of parthenogenesis costs i.e *C*_*p*_ = 0, the condition in which a high fertilization rate is the only evolutionary outcome is 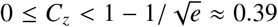(see Figure 3). Since situa-tions in which the fertilization rate initially tends to zero are not of interest in the current analysis, we work in the remainder of this paper in the regime in which increasing fertilization rate is selected for; that is, 1 ⪆ *C*_*z*_ and with initial conditions that do not lead to a stable *α* = 0 fixed point (see Figure 3, unshaded regions).

### 3.2. Evolutionary branching can lead to anisogamy, followed by “oogamy” (or suppression of pheromone production)

In Figure 3 we see that the approximation obtained for the co-evolutionary dynamics of *m* and *α* in a single fixed environment, Eq. (5), accurately captures the dynamics of the full model realised via numerical simulation at early times. One point of departure is that at long times as *α* increases along the *m* ≈ *β*/4 manifold, we see the mean mass trait value from simulations increasing to higher values than those predicted analytically. In Figure 4, we show that this is a result of evolutionary branching in the simulations.

**Figure 4:**
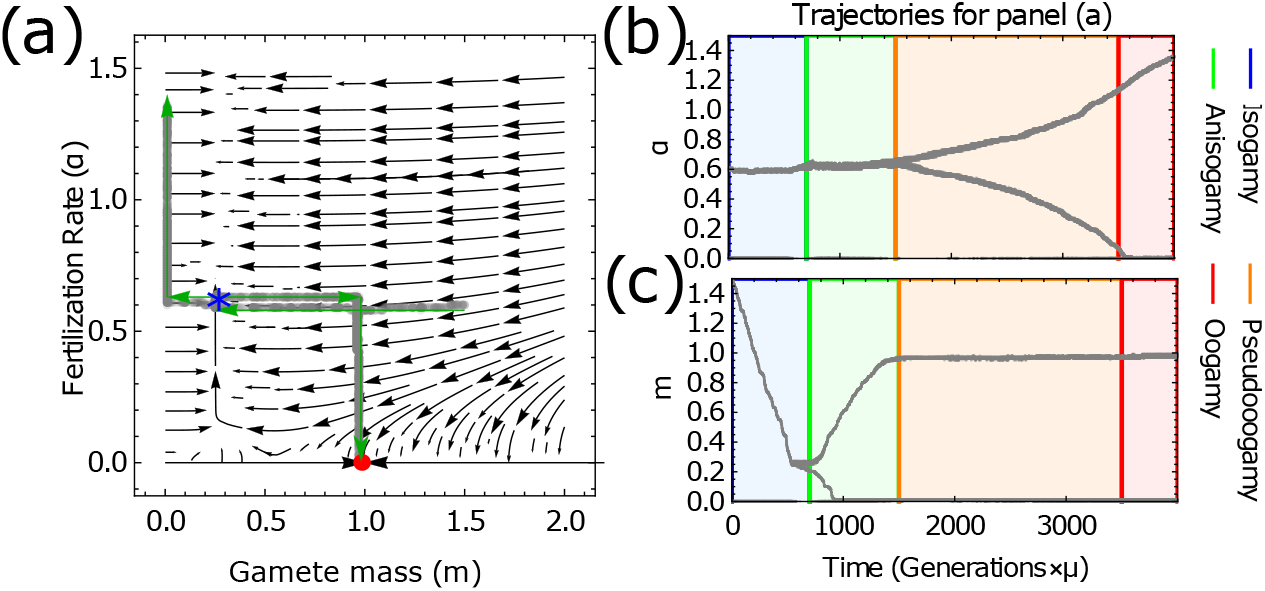
Numerical illustration of evolutionary branching in Figure 3(b). Panel (a): Analytic predictions for the early evolutionary dynamics (as Figure 3(b)) overlaid with trajectories (*m*_*i*_(*t*), α_*i*_(*t*))) for each *i*^th^ trait. Evolutionary branching is observed along the *m* ≈ *β*/4 manifold, indicated by the blue star. Green arrows show the temporal progression of the branching. Panels (b) and (c): The temporal trajectories of the traits α_*i*_(*t*) and *m*_*i*_(*t*) respectively, showing that the evolutionary trajectory passes from isogamy to oogamy. The starting point of the trajectory is (*m*(0), α(0)) = (1.5, 0.6). Parameters used are *A* = 100, *M* = 1, *T* = 1, *C*_*z*_ = 0.6, *C*_*p*_ = 0, *β* = 1, *δ* = 0.01, µ = 10^−3^, *f*_0_ = 2 × 10^−3^ and simulation run for 3.5 × 10^6^ generations.

Should the evolutionary trajectories reach a sufficiently large fertilization rate, *α*, along the *m* ≈ *β*/4 manifold, anisogamy can evolve through evolutionary branching in mass (see Figure 4), which gives rise to a dimorphic gamete population that contains one large gamete (macrogamete) and one small gamete (microgamete) with masses

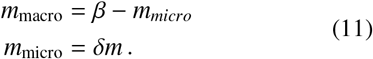

Here the microgamete evolves to the smallest possible nonzero trait value in our simulation model (*δm*) while the macrogamete evolves to produce macrogamete-microgamete zygotes of mass *β* (see Appendix G for simulation). Analytically, this branching is shown to be due to disruptive selection in mass in the vicinity of the *m* ≈ *β*/4 manifold at high values of *α*. In Appendix H, we show that this region is an approximate evolutionary singularity. This initial branching has been noted in other models that also do not account for self-incompatible mating types but that consider the case of fixed *α* and obligate sexual reproduction [24].

As the mass of the macrogamete becomes larger (and that of the microgamete smaller), a branching in fertilization rates can also occur (see Figure 4 (b)). Near this mass *β* − *m*_*micro*_, the macrogamete has a high survival rate under parthenogenesis, and the benefits of fertilization (in particular with microgametes of very small mass) can be outweighed by the costs of fertilization. Selection can thus act to lower the fertilization rate of macrogametes, *α*_macro_, towards zero. This in turn leads to an increased selection pressure for the microgametes to increase their fertilization rate, with *α*_micro_ → *α*_max_, to increase the probability of microgametes fertilizing macrogametes (averting the low survival probabilities of microgametes under parthenogenesis).

The situation described above is one in which macrogametes are still fertilized by microgametes (at a rate *α*_micro_/2) but do not fertilize themselves. Biologically, one interpretation of this situation is the evolution of oogamy, with *α*_micro_ → *α*_max_ and obligate sexual reproduction in the limit *α*_max_ → ∞. However, since *α* is a compound parameter the captures both motility and other mechanisms to enhance cell encounter rate, this situation can also be viewed as one in which macrogametes have reduced pheromone production, gamete recognition protein affinities, or any other mechanism to limit fertilization by microgametes. For concision however, we will refer to this state simply as oogamy for the remainder of the results section.

Above we have shown that when costs to fertilization are accounted for, a continuous evolutionary trajectory can exist that takes the population from a state of isogamy to oogamy. In the following sections we demonstrate how each of these transitions can be arrested under various parameter regimes.

### 3.3. A low ceiling on the fertilization rate can stabilize isogamy

If the maximum possible fertilization rate *α*_max_ is limited to a low value, for instance due to energetic constraints on motility or environmental constraints such as turbulence, then it is possible to prevent the transition from isogamy to anisogamy. In Appendix H, we derive the minimum value of *α*_*max*_ at which evolutionary branching in mass can occur, and find that if

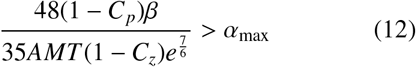

evolutionary branching in mass cannot occur. These results are illustrated in Figure 5. Should *α*_max_ lie below this threshold (equivalent in approximately 70% of cells gametes fertilized in Figure 5, see Figure B.10) the population is held in a state of isogamy.

**Figure 5:**
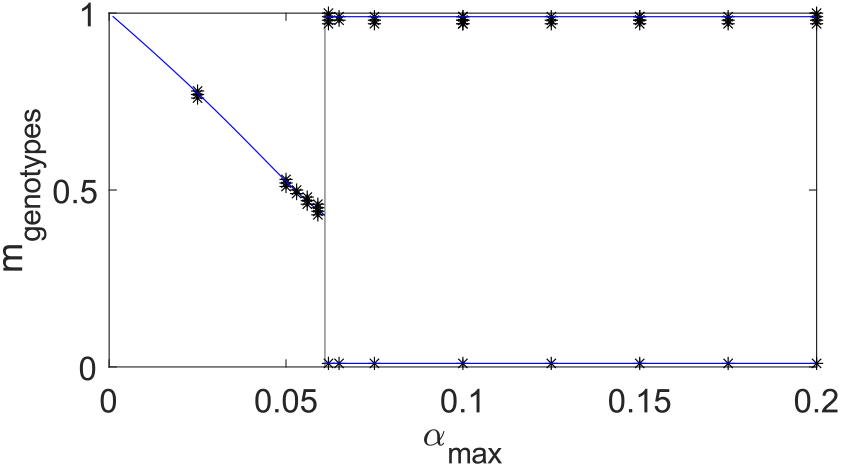
Analytical prediction (blue) and numerical illustration of the range of *α*_*max*_ in which branching to anisogamy is possible. Black markers represent the mass of each gamete genotype within the population after 2.5 × 10^6^ generations. Once branching to anisogamy has occurred, a dimorphic gamete population, characterised by the presence of two genotypes would be present. One genotype where *m* ≈ *β* − *δm* and one where *m* = *δm*. Vertical line represents the analytically predicted *α*_*max*_ above which branching can occur in *m* Eq. (12). The blue curve left of this line is the numerical solution to 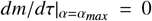 in Eq. (E.1) and the blue lines towards the right contains the theoretical masses of the macrogamete (*m* = *β* − *δm*) and microgametes (*m* = *δm*). Parameters are *A* = 100, *M* = 1, *T* = 0.1, *C*_*z*_ = 0.3, *C*_*p*_ = 0, *β* = 1, *δ* = 0.01, µ = 10^−3^ and *f*_0_ = 2 × 10^−3^.

### 3.4. High costs of parthenogenesis relative to zygote formation can stabilize anisogamy (or promote macrogamete pheromone production)

Analytically, we show in Appendix I that when costs for zygote formation are relatively low and costs for parthenogenesis relatively high, a fully “oogamous” state (in which the macrogamete’s fertilization rate evolves towards zero, *α*_macro_ → 0) does not evolve following the transition to anisogamy. The costs to fertilization must instead be sufficiently high, such that

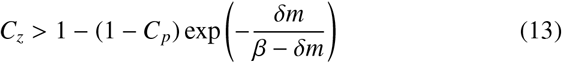

for true oogamy to evolve. This is illustrated in Figure 6. Essentially if the costs to the macrogamete of fertilization are not sufficiently high, then there is no longer a selective pressure (as described in Section 3.2) for the macrogamete to avoid fertilizations by decreasing its fertilization rate. While the population does not evolve towards oogamy, its ultimate state is dependent on the maximum fertilization rate, *α*_max_.

**Figure 6:**
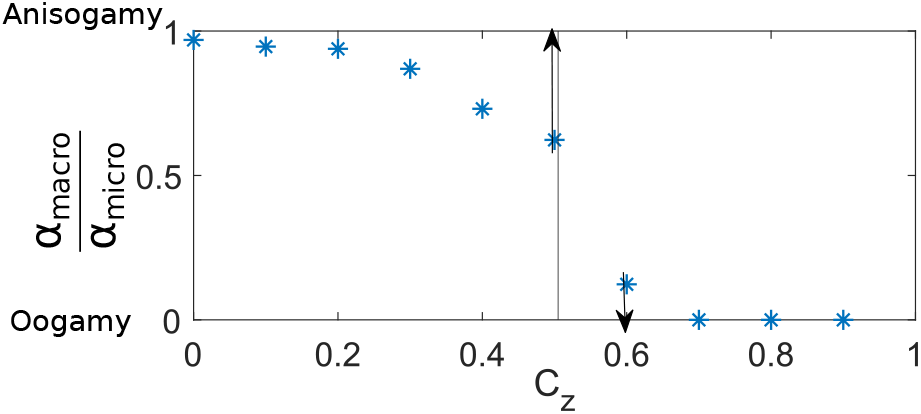
Numerical illustration of the ratio of macrogamete to microgamete fertilization rate, *α*_*macro*_/*α*_*micro*_. Oogamy is favoured over anisogamy above a sufficiently high fertilization cost *C*_*z*_, predicted analytically by the vertical line (see Eq. (13)). Here *C*_*p*_ = 0.5, (*m*(0), *α*(0)) = (0.25, 0.02) and *α*_*max*_ = 1.3. All other parameters are as in Figure 3 except *f*_0_ = 0.02 and the simulation is run for 6 × 10^6^ generations. Back arrows point in the direction towards which *α*_*macro*_/*α*_*micro*_ evolves in infinite time.

Suppose we initially place no limit on the maximum fertilization rate, *α*_max_ → ∞, and that the costs *C*_*p*_ and *C*_*z*_ are such that there is no longer selection for the fertilization rate of the macrogamete to decrease (i.e. the inequality in Eq. (13) does not hold). While there may be a selective pressure for the fertilization rate of the macrogamete to increase, this selection pressure is weaker for the parthenogenetic macrogamete than the microgamete, as the microgamete relies more heavily on the fertilization pathway for its survival. Thus although the fertilization rates of both macrogametes and microgametes, *α*_macro_ and *α*_micro_, evolve to increase *α* indefinitely, the macrogamete does so at a considerably slower rate. Under this scenario, we therefore observe pseudooogamy [27], where the macrogamete fertilization rate does not evolve toward zero but is still lower than the microgamete fertilization rate. Equivalently, we expect to see parthenogenetic macrogametes invest less in mechanisms such as pheromone production that increase fertilization rate than parthenogenetic microgametes. While this situation is only observed when *α*_max_ → ∞, in the case of large but finite *α*_max_, pseudooogamy is a prolonged transient state (see Figures 6 and I.19). Lastly, should *α*_max_ be smaller, such that the increasing fertilization rate of macrogametes can “catch up” with that of microgametes, the population can return to anisogamy following a transient period of oogamy.

### 3.5. In a switching environment, the population can evolve a bet-hedging strategy that stabilizes isogamy

In Figure 7, we see that the analysis for the evolutionary dynamics in the case of a switching environment (see Eq. (9)), provides a good approximation to the dynamics of the full model (which accounts for multiple traits coexisting under a mutation-selection balance) realised via numerical simulation. We now see three broad evolutionary outcomes.

**Figure 7:**
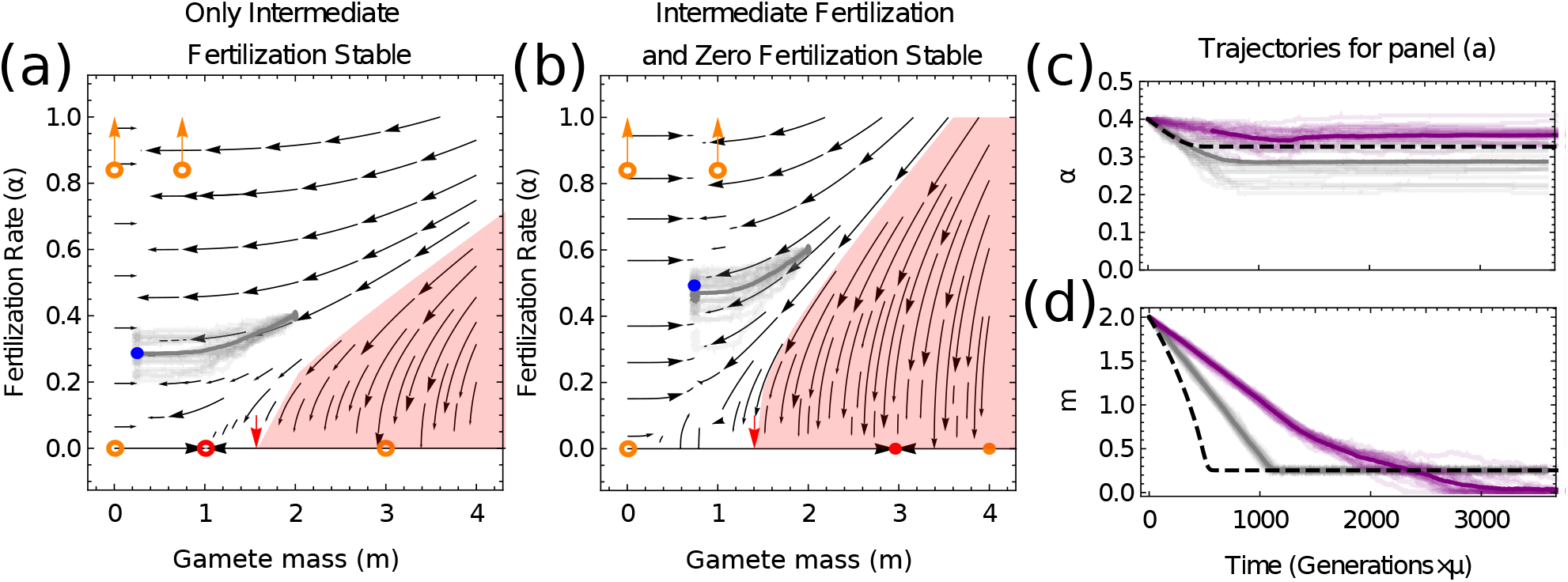
Phase portraits for the co-evolutionary dynamics in a switching environment (see Eq. (9)). In addition to qualitatively similar dynamics as in the fixed environment (see Figure 3), two new evolutionary scenarios are now possible, including populations in which stable intermediate fertilization rates (filled blue circle) are the only evolutionarily stable state (see panel (a)) and populations in which there is an additional zero fertilization stable state (filled red circle, see panel (b)). Open orange circles represent the (now unstable) states to which the population can be attracted in either environment 1 or 2 (where *β* = *β*_1_ or *β* = *β*_2_). Average population trait trajectories, (⟨*m*⟩ (*t*), ⟨*α*⟩ (*t*)), from simulation of the full stochastic model in the FRTI regime are overlaid in gray. In panel (c) we plot the time-series in the FRTI regime (gray, as in panel (a)) alongside those in the FRTE regime (purple) and our analytic predictions (black-dashed). In all panels, *A* = 100, *M* = 10, *T* = 0.1 and *f*_0_ = 2 × 10^−3^. Under the FRTI regime (overlaid in gray in panels (a), (b) and (c)), µ = 3.5 × 10^−4^, *δ* = 7 × 10^−3^ and the simulation is run for 10^7^ generations. Under the FRTE regime (overlaid in purple in panel (c)), µ = 5 × 10^−3^, *δ* = 5 × 10^−3^ and the simulation is run for 7 × 10^6^ generations. In panel (a) (and panel (c) gray), *C*_*z*_ = 0.35, *β*_1_ = 3, *P*_1_ = 0.335, (*m*(0), *α*(0)) = (2, 0.4), λ_1→2_ = 0.250 and λ_2→1_ = 0.126. In panel (b), *C*_*z*_ = 0.7, *β*_1_ = 4, *P*_1_ = 0.74, (*m*(0), *α*(0)) = (2, 0.6), λ_1→2_ = 5.86 × 0.01 and λ_2→1_ = 0.167. In panels (c) and (d), the switching rates for the FRTE regime (purple) are λ_1→2_ = 2.93 × 10^−5^ and λ_2→1_ = 8.34 × 10^−5^. Switching rates and mutation rates µ are all measured in units of (number of generations)^−1^ and *C*_*p*_ = 0 in all panels.

We begin by considering the intuitive limit of *τ*_1_ ≫ *τ*_2_. In this scenario the population spends almost all of the time in environment 1, and a comparatively insignificant amount of time in environment 2 (i.e. *P*_1_≈ 1 and *P*_2_ ≈ 0). Consequently, the population evolves approximately as if it were simply in a fixed environment with *β* = *β*_1_, and the conditions given for the fixed environment, Eq. (10), can be used to infer the evolutionary outcome. An analogous argument holds for the dynamics when *τ*_2_ ≫ *τ*_1_, but with *β* = *β*_2_ in Eq. (10). Once on the *m* = *β*_*i*_/4 manifold towards the high-fertilization rate fixed point, evolutionary branching towards anisogamy and oogamy can occur as described in Section 3.2. When the time spent in each environment is of the a comparable order however (e.g when *τ*_1_, and *τ*_2_ have not entirely dissimilar magnitudes), we find the emergence of conservative bet-hedging strategies (see Figures 7, 8 and Eq. (16)) where the population evolves to produce a single genotype adapted moderately to both environments [72, 74].

**Figure 8:**
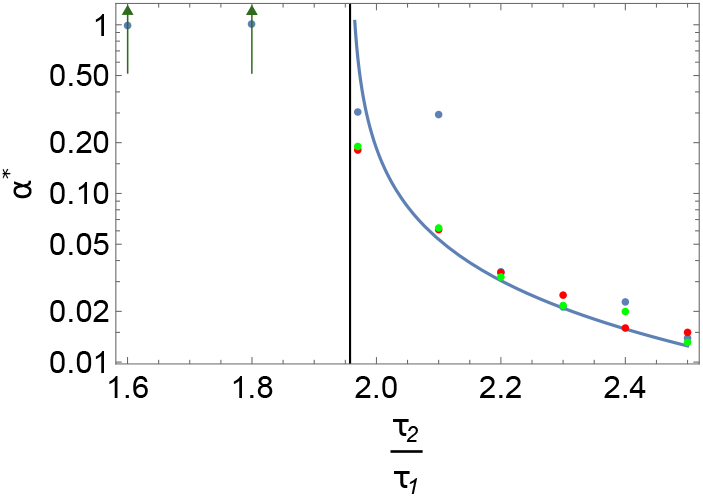
Analytic theory for the fertilization rate at the switching-induced isogamous fixed point (blue line, see Eq. (16)) against simulations (dots). The black vertical line is the analytic prediction for when this fixed point vanishes, with parameters left of the line corresponding to destabilized isogamy. Simulations are obtained by averaging over 25 realisations of ⟨*α*(*t*)⟩ corresponding to stochastic evolutionary trajectories. Parameters are *δ* = 0.01, µ = 5 × 10^−4^, *f*_0_ = 2 10^−3^, τ_1_ = 0.25 and simulation run for 1.2 × 10^7^ generations. All other parameters are the same as in the FRTI regime of Figure 7 (a). Red markers are for (*m*(0), *α*(0)) = (0.29, 0.075), green for (*m*(0), *α*(0)) = (0.29, 0.15) and blue for (*m*(0), *α*(0)) = (0.29, 0.3).

Under a range of parameter conditions, we find analogous evolutionary attractors for the fertilization rate as in the fixed-environment case (see Eq. (10) and Figure 7), but with bet-hedging strategies [68] for the gamete mass;

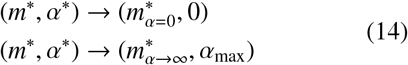

with

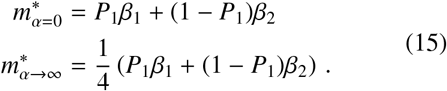

The population thus initially evolves either to high fertilization rates or zero fertilization rates, but with a mass which is the weighted average of the optimal strategy in either environment.

Under a more restricted set of parameter conditions however we find that a bet-hedging strategy for the fertilization rate can also evolve; essentially a switching-induced fixed point can manifest, as illustrated in Figures 7 and 8. Here the tension between the evolutionary dynamics in the two environments (which can select for high fertilization rate and large gametes in one environment, and zero fertilization rate and small gametes in the other) can lead to the population being held in a state at which intermediate finite fertilization rate and isogamy form the evolutionarily stable bet-hedging strategy. In Appendix F.3, we show that the switching-induced fixed point is given by

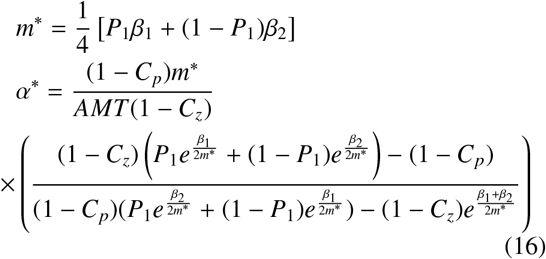

In Figure 7(c-d) we see that this switching induced fixed point is observed in the evolutionary simulations in which multiple traits can be held in the population under a mutation-selection balance. In the FRTI regime (gray lines), in which the environment changes more quickly, the population is held at the predicted mass *m*, while more variability between simulations is seen around the predicted fertilization rate *α* as a result of the weaker selection on this trait. In the slower-switching FRTE regime (purple lines), a greater quantitative difference between the results of simulations and the analytic predictions is observed. However in both regimes, the key prediction of finite fertilization rate (Figures 7 and 8) and isogamy is indeed captured (see Figures F.15 and F.16).

## 4. Discussion

In this paper we have extended classic results on the evolution of anisogamy [14, 25] to account for parthenogenetic development [41, 42], the co-evolution of fertilization rate and gamete cell mass, and stochastically varying environments. In doing so we have demonstrated the possibility of a continuous evolutionary trajectory from an initial state of isogamy to anisogamy followed by oogamy, an evolutionary trajectory observed throughout the eukaryotes [3, 10].

Consistent with earlier theoretical results on the evolution of anisogamy that neglected the pre-existence of self-incompatible mating types [24], we have shown that anisogamy can evolve from isogamy through evolutionary branching. However our model also shows that when the possibility of parthenogenetic development is accounted for, a subsequent branching is possible, comparable to oogamy. Importantly, this transition is possible even without explicitly accounting for pheromone-receptor systems [29, 30], speed-size relationships [32, 34], mechanistic gamete encounter rates [31] or costly motility [75], as in previous work. Amongst these, the model we present here most closely resembles that of [34], which explored the coevolution of mass and investment in motility to study how investment in mating differed between the two sexes. Our fertilization rate parameter *α* can be thought of as a crude compound parameter that captures the effect of motility, surface area for collision and the collision probability as entailed in [34]. However unlike previous work that does not consider the possibility of parthenogenesis, the fertilization rate parameter *α* also captures the effect of gamete recognition protein affinity.

Importantly, while selection for oogamy or pseudooogamy in earlier work is driven by selection to increase fertilization rates in species with obligate sexual reproduction [29, 30, 32, 34, 31], in our model selection for oogamy is driven by sexual conflict [76] and a selection pressure for parthenogenetic macrogametes to *reduce* their fertilization rate. In particular, microgametes receive a survival advantage by increasing their fertilization rate with macrogametes, while macrogametes evolve to zero fertilization rate to overcome fertilization costs. This requirement of fertilization costs for oogamy to evolve under parthenogenesis provides a complementary result to [27], where instead high motility costs were required for oogamy.

Given the selection pressure for macrogametes to avoid fertilisation in these costly scenarios, it is natural to ask why macrogametes should not evolve defensive traits to prevent fertilisation by microgametes. Such traits are possible; selection on eggs to reduce the affinity of their gamete recognition proteins is understood to operate under high sperm competition (high sperm density) in order to reduce the probability of detrimental polyspermy [49] or to select for fit sperm/paternal genotypes, or to limit polyspermy [77]. A simple answer is that we have not accounted for the genetic benefits provided by sex [78], which would drive additional selection for fertilisation. However deeper insights can be developed by considering the empirical literature.

The eggs of parthenogenetic brown algal species are known to produce pheromones in general [79]. This is also true of the brown alga *Scytosiphon lomentaria* in the southern seas around Japan, where female gametes release pheromones to attract male gametes [80]. However, in the northern seas around Japan, female *S. lomentaria* gametes can exhibit completely suppressed pheromone production and populations consist entirely of females reproducing parthenogenetically [80]. These empirical results build on earlier work showing that females from the parthenogenetic populations release gametes that are larger and produce lower levels of sex pheromones [81]. Interestingly, these qualitative behaviours are broadly captured by our model if one assumes that colder sea temperatures can be interpreted as an increase in the environmental harshness parameter, *β*, as illustrated in Figure 9. Essentially, increasing *β* (as might be expected in colder waters) can lead to Eq. (13) being satisfied such that *α*_*macro*_ → 0 (e.g. pheromone production is switched off) while simultaneously leading to increased macrogamete size according to Eq. (11).

**Figure 9:**
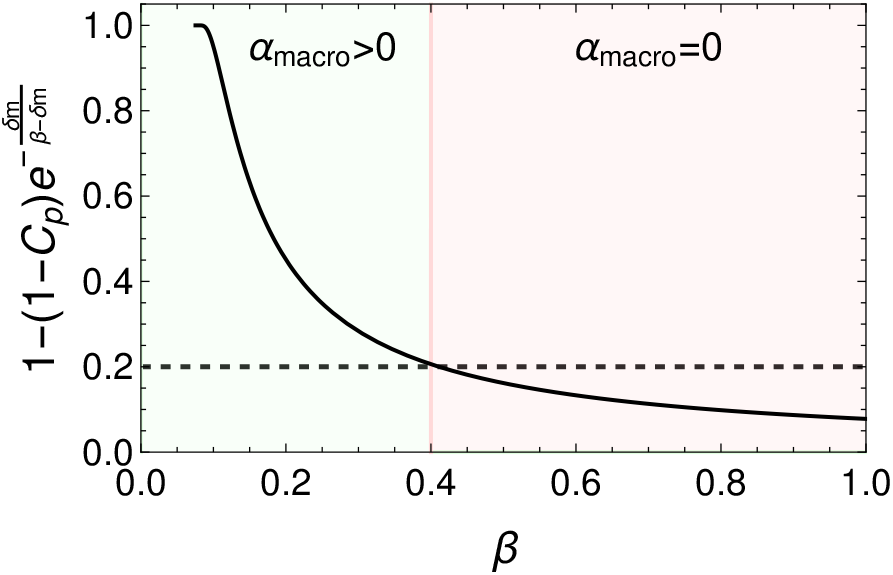
Plot illustrating the suppression of macrogamete fertilization when *β* becomes large. Black line shows the right hand side of the inequality in Eq. (13). When this calls below *C*_*z*_ (black dashed line) selection acts to drive the macrogamete fertilization rate *α*_max_ to zero. Thus the model predicts that as the harshness of the environment *β* increases, macrogametes should evolve to resist fertilization and grow larger (see Eq. (11)). Parameters used are *C*_*z*_ = 0.2, *C*_*p*_ = 0 and *δm* = 0.075.

We have shown analytically that various conditions exist that can arrest the population at different stages of the isogamy-anisogamy-oogamy evolutionary trajectory. In particular our model suggests that isogamy can be stabilized under greater energetic constraints or in highly turbulent environments. We have also shown that isogamy can be stabilized as a bet-hedging strategy under switching environmental conditions [68, 72]. Here, in organisms that reproduce parthenogenetically, microgametes that fail to find a partner face low survival prospects under harsh environmental conditions, and thus the transition to anisogamy is frustrated. Such dynamics may be at play in isogamous species that reproduce parthenogenetically, such as those among the *ectocarpus* [82] and *Blidingia minima* [37].

The results described above are dependent on the capacity for parthenogenetic development amongst gametes that fail to find a partner to fertilize with. Such a capacity is widespread in the brown algae [51] and can also be found in the green algae [41]. We have also shown that for the evolution of oogamy in our model, we require costs of fertilization to exceed those of parthenogenetic development. While empirically parthenosporophytes have lower survival probabilities than zygotes [41], disentangling the costs of development pathways (parthenogenesis or fertilization) from those of increased mass is challenging [83, 42]. By invoking costs to zygote formation arising from cytoplasmic conflict [58], our work is reminiscent of alternative hypotheses for the evolution of anisogamy [59, 60], that suggest that anisogamy reduces the potential for such conflict by limiting cytoplasmic contributions from the microgamete. However many other costs to fertilization could be present, including the possibility of failure during cell fusion [57], and the energetic and time costs of sexual reproduction [62]. Unpacking and quantifying these costs remains an interesting area of empirical research [79].

Our primary reason for neglecting the existence of mating types in our model was for mathematical simplicity [24]. Including such types in a model would be a natural next step, and based on the biological reasoning above, such a model should demonstrate similar qualitative behaviours. However it is worthwhile noting that many models of the evolution of sexual reproduction in early eukaryotes suppose the existence of a “uni-sexual” early ancestor that mated indiscriminately [84]. Thus our model, which has neglected the existence of mating types, could be seen as reflecting such a system. In this context, the evolution of incompatibility between macrogametes in our model (that fuse with each other at rate zero in the oogamous scenario) is interesting, as the origins of self-incompatible mating types are still debated [85, 86]. This mechanism would represent an “anisogamy consequence” model for the evolution of mating types that manages to identify the conditions under which fusion between large macrogametes is disadvantageous [87]. However as such “anisogamy consequence” models are not consistent with most empirical observations (that suggest mating types preceded anisogamy), this interpretation should be treated with caution.

As with any model, there are omissions from our formulation. These include the mortality of gametes (known to stabilize isogamy [25]) the discrete nature of cell divisions leading to gametes [41], and non-local trait mutations. While these would be interesting additions to our model, the key insights derived from the PBS model have remained remarkably robust to such generalizations, and so the inclusion of these additional considerations to our model may lead primarily to quantitative, rather than qualitative, changes in results. More notably, as motility and area of collision can be influenced by mass, it will be important to account for mass dependencies in future. As shown in [34], under a positive speed-size relation, mass and fertilisation rate are likely to both evolve to increase, while under a negative speed-size relation, gametes that evolve a high motility would be expected to evolve small masses. Similarly, costs to sexual reproduction include pheromone production. Such costs may therefore be a function of the fertilization rate, which we have not accounted for here. More generally, extending our mathematical approach leveraging adaptive dynamics to switching environments [68] in other facultatively sexual populations might prove particularly fruitful [88, 89].

In this paper, we have extended the models of [14, 24] in several ways; by allowing the fertilization rate to evolve, accounting for the possibility for unfertilized gametes to develop parthenogenetically should they fail to locate a partner, and subjecting the system to switching environments. In doing so, we have shown its capacity to parsimoniously capture continuous evolutionary trajectories from isogamy to oogamy in parthenogens, as well as the suppression of pheromone production in parthenogenetic external fertilizers. Moreover, our models emphasise the importance of investigating the co-evolutionary dynamics for a range of evolutionary parameters and their implications for the evolution of fertilization rates in parthenogens.

## Appendix A.

### Individual sporophytes must produce twice as many spores as Parthenosporophytes to maintain the same reproductive output

Here, we show that diploid sporophytes must produce twice as many meio-spores as are produced by haploid parthenosoprophytes in order to maintain a parity in reproductive output (see Figure 2). Further, we show that meio-spore production in sporophytes below this level constitutes an implicit cost of fertilization.

Recall that, in the main text, *C*_*z*_ is the survival cost incurred by taking the sexual reproductive route while *C*_*p*_ is the survival cost incurred by taking the parthenogenetic reproductive route. We denote *F*(*T*) as the number of fertilized zygotes, 2*F*(*T*) as the number of haploid gametes that have formed zygotes at the end of a generation, and *N*(*T*) as the number of unfertilized gametes at the end of a generation. The functional form for the absolute fitness of a single genotype is

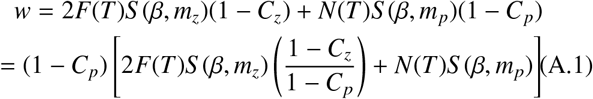

where *m*_*z*_ is the mass of fertilized zygotes, *m*_*p*_ is the mass of unfertilized gametes and *S* (*β, m*) is the survival function (see Eq. (2)).

Alternatively, we can express the model in terms of the number of sporophytes and parthenosporophytes, and their individual rates of meio-spore/spore production, *G*_*z*_ and *G*_*p*_ respectively. We again denote the number of zygotes (destined to become diploid sporophytes) as *F*(*T*) and the number of unfertilized gametes (destined to become haploid parthenosporophytes) as *N*(*T*). The absolute fitness of a single genotype can now be written

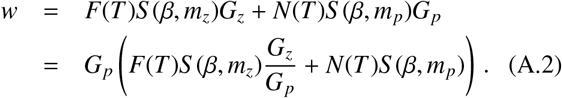

We now note that in terms of relative fitness, the constant factors (1 − *C*_*p*_) and *G*_*p*_ preceding Eqs. (A.1-A.2) are inconsequential.

Now equating the pre-factors of *F*(*T*)*S* (*β, m*_*z*_) in Eqs. (A.1-A.2) allows us to evaluate the mass-independent costs to zygote formation in the model. We see that

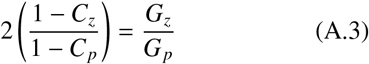

which we can rearrange in terms of *C*_*z*_ to get

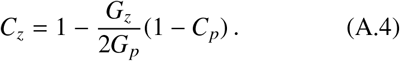

For zero costs to parthenogenesis, *C*_*p*_ = 0, production of meio-spores by sporophytes (*G*_*z*_) must be twice that of spores by parthenosporophytes (*G*_*p*_) in order to achieve zero cost to zygote formation. This result has a straightforward biological interpretation. Since under Mendelian inheritance the reproductive fitness of sporophytes is shared between the gametes that contribute towards its production, the fitness of sporophytes must be at least twice that of parthenosporophytes to avoid an implicit cost to zygote formation.

## Appendix B. Within generation dynamics

At the start of each generation, we assume a total of *A* organisms of mature cell size (or energy budget) *M* divide to form gametes. We further assume that the frequency of mutants in this adult population is given by 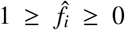, with *i* = *m* for mutants that change the mass of gametes and *i* = *α* for mutants that change the fertilization rate of gametes.

### Appendix B.1. Fertilization kinetics: mutant with different mass

If a mutation occurs changing the size of gametes produced by the mutant (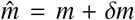 for resident gamete mass *m*), the fertilization kinetics themselves (see Eq. (1)) are unaffected by the change in mass. However the number of gametes produced by the mutant will change. Denoting by *N* the number of resident gametes and 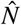 the number of mutant gametes, we have

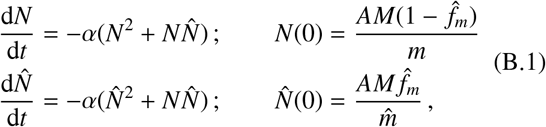

which has a solution

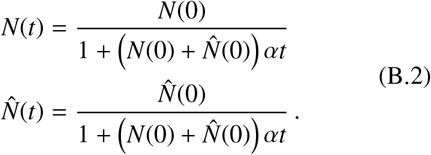

This allows us to determine the number of unfertilized cells of each type at the end of the fertilization window at *U* = *N*(*T*) and 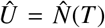, where we recall that *T* is the length of the fertilization window.

We also need to determine the number of fertilized cells formed from the fertilization of two residents, 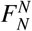, two mutants, 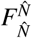, and a mutant and a resident, 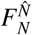. For this we need to solve

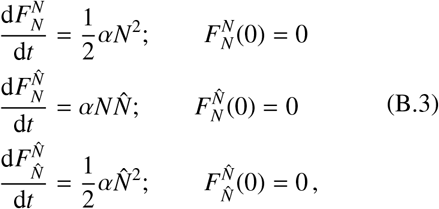

which can be solved by substituting for *N* and 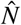 from Eq. (B.2) to yield

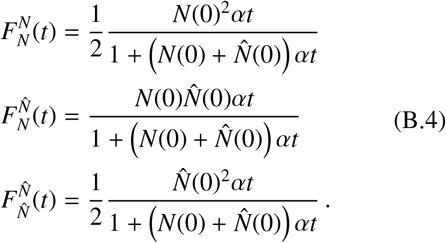

### Appendix B.2. Fertilization kinetics: mutant with different fertilization rate

If a mutation occurs changing the fertilization rate of gametes produced by the mutant (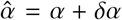 for residents with fertilization rate *α*), the fertilization kinetics themselves are altered relative to Eq. (1). We assume for simplicity that the fertilization rate between resident-pairs is the mean of the fertilization rate between the two types in isolation, such that

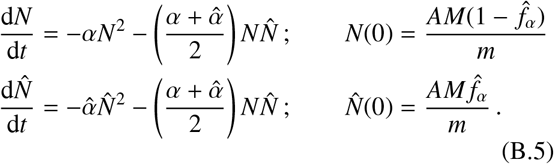

Solving this equation is slightly less straightforward than solving Eq. (B.1). However we can make analytic progress by making a change of variables and applying an approximation based on small mutational step size *δα*.

We introduce the transformed variables 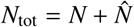 and 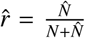, representing the total number of unfertilized cells and the frequency of unfertilized mutant cells respectively. Eq. (B.5) then becomes

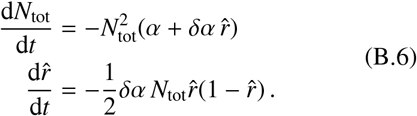

We now see that although this equation is also intractable, the leading order dynamics of *N*_tot_ are governed by *α*. Therefore when *α* ≫ *δα*, we make the approximation 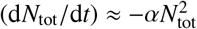 We then obtain

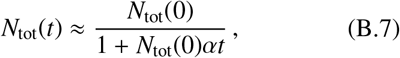

and substituting this into our equation for 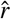 in Eq. (B.6), we can solve to obtain

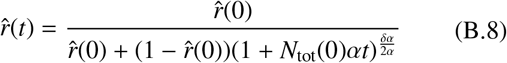

Inverting the transformation, we then arrive at

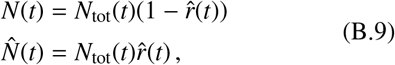

with *N*_tot_ and 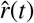 taken from Eqs. (B.7-B.8). We shall see in Appendix B.4 that calculating 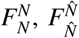, and 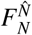 explicitly is in fact unnecessary, and these expressions for *N*(*t*) and 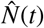 are sufficient for analytical progress.

### Appendix B.3. Change in mutant frequency over a generation: mutant with different mass

We begin by calculating the fitness of the resident and a mutant that changes the mass of gametes, *w*_*m*_ and 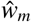 respectively, which are simply given by the total number of cells of each type at the end of a generation. Recalling that fertilized and unfertilized gametes both survive with a probability governed by the parameter *β* and the mass of the cell *m*_*c*_ (see Eq. (2)), and that fertilized cells survive with an additional probability (1 − *C*_*z*_), we have

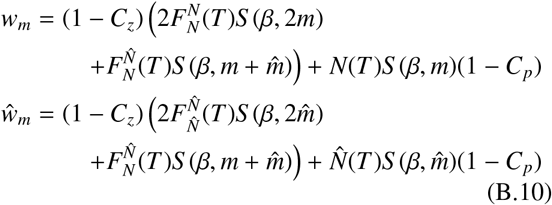

Where 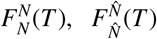, and 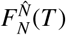 are taken from Eq. (B.4), and *N*(*t*) and 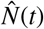 are taken from Eq. (B.2). These expressions can be used to calculate the frequency at the end of the generation, 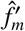, of a mutant that changes the mass of gametes as

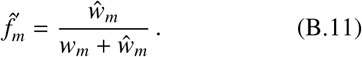

The change is the frequency of the mutant over the course of a generation is then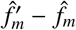.

**Figure B.10:**
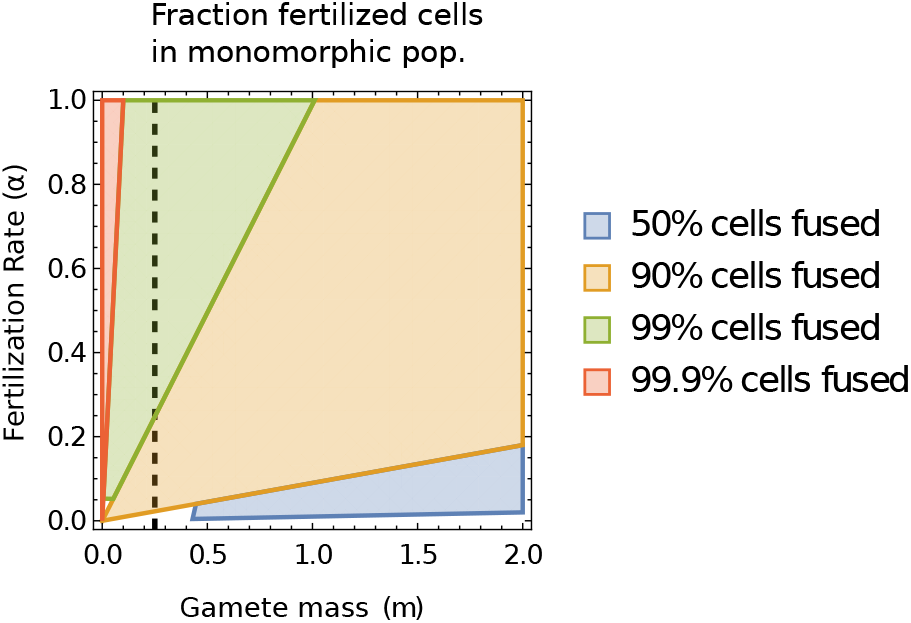
Illustration of the total proportion of cells that are fertilized at the end of a fertilization period (length *T* = 1) in a monomorphic isogamous population (without branching) as a function of trait variables *m* and *α*. Parameters used are the same as those in Figure 3. The vertical black dashed line gives the location of the manifold (*β*/4, *α*), along which the population is attracted to when approaching the high *α* fixed point.

### Appendix B.4. Change in mutant frequency over a generation: mutant with different fertilization rate

Taking an analogous approach to Appendix B.3, we begin by calculating the fitness of the resident and a mutant that changes the fertilization rate, *w*_*α*_ and 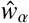 respectively. We obtain

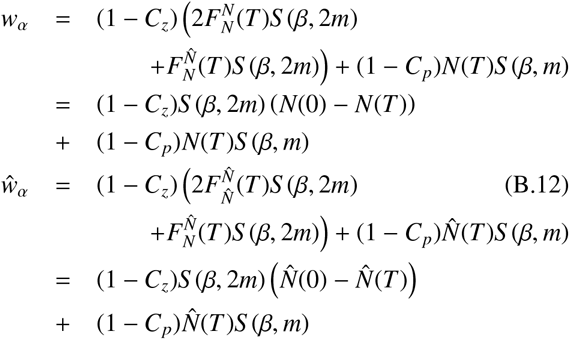

where *N*(*t*) and 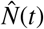 are now taken from Eq. (B.9). Here we have used the fact that since the survival function for fertilized cells, *S* (*β*, 2*m*), is independent of the composition of the fertilized cells (mutations here only affect fertilization rate) the number of cell-types contributing to the fertilized cells can be inferred under cell conservation during the fertilization period (e.g. for resident cell types 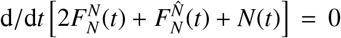, and similarly for mutant cell types).

These expressions can be used to calculate the frequency at the end of the generation, 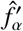, of a mutant that changes the fertilization rate as

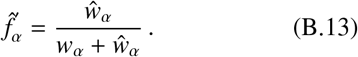

The change is the frequency of the mutant over the course of a generation is then 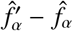.

## Appendix C. Invasion dynamics

Our aim is to derive the dynamics of the frequency of a mutant with a small mutation in either of the traits, *m* or *α*, over multiple generations. We introduce *t*_*g*_ as a measure of the number of discrete generations.

### Appendix C.1. Invasion ODE: mutant with different mass

We begin by deriving the dynamics for the frequency of a mutant that changes the mass of gametes. We begin by assuming that the mutational step size, *δm*, is small. Under these conditions, the frequency of mutants changes only by a small amount over the course of one generation, and we can approximate the frequency of mutants at the end of the generation (see Appendix B.3) by 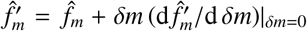 where 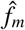 is the frequency of the mutants at the beginning of the generation. The dynamics of the mutant frequency over an invasion can then be approximated by

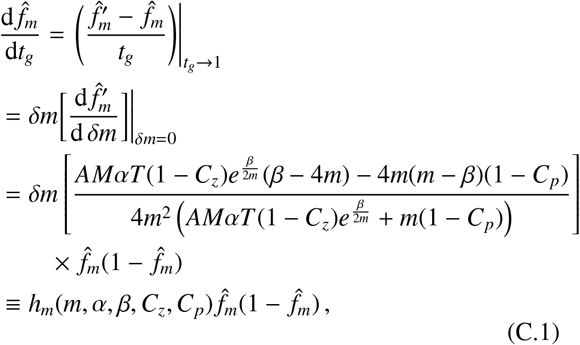

(see also Eq. (3)), where 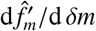 is derived from Eq. (B.11). We show in Figure C.11 that as expected, this is a good approximation for the dynamics when *δm* is small.

### Appendix C.2. Invasion ODE: mutant with different fertilization rate

We now derive the dynamics for the frequency of a mutant that changes the fertilization rate of gametes. Taking an analogous approach to Appendix C.1, we assume the mutational step size for mutation, *δα*, is small. We can then approximate the frequency of mutants at the end of the generation (see Appendix B.4) by 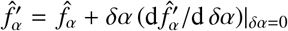, where 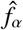 is the frequency of the mutants at the beginning of the generation. The dynamics of the mutant frequency over an invasion can then be approximated by

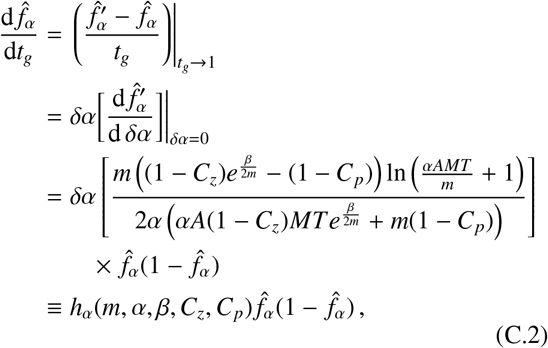

(see also Eq. (4)) where d 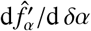 is derived from Eq. (B.13). We show in Figure C.12 that as expected, this is a good approximation for the dynamics when *δα* is small.

**Figure C.11:**
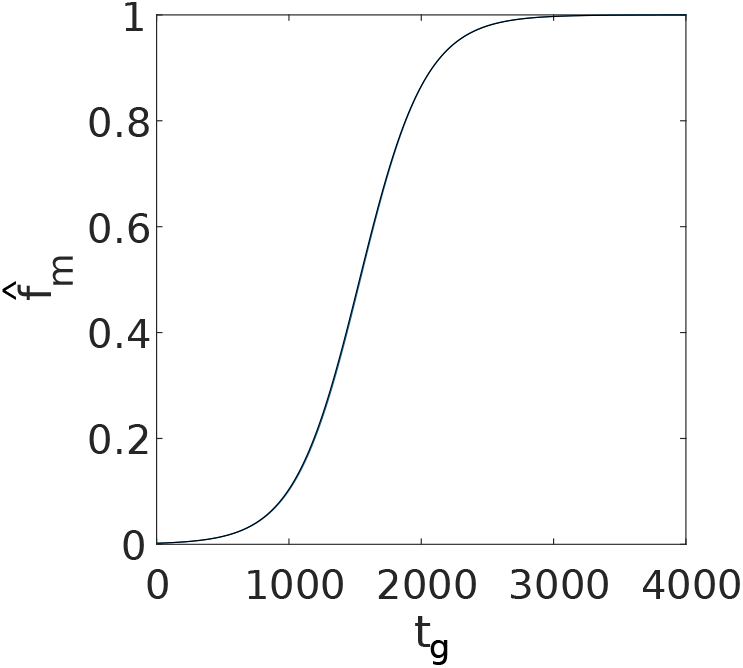
Invasion dynamics for a mutant with mass *m* + *δm*. Blue - analytical prediction using Eq. (C.1), black - numerical simulation. The initial condition is (*m*(0), *α*(0)) = (0.4, 0.1). Parameters are *δm* = − 0.005, *f*_0_ = 0.002, *G* = 4 ×10^3^, *A* = 100, *M* = 1, *T* = 1, *C*_*z*_ = 0.6, *C*_*p*_ = 0 and *β* = 1.

## Appendix D. Deriving Evolutionary dynamics using a multidimensional approach

The evolutionary dynamics can be derived without specifying the invasion dynamics using the standard multidimensional approach of evolutionary analysis given in [90, 91, 92] (which does not require the full derivation of the invasion trajectories calculated in Appendix C). First, we consider the fertilisation kinetics, which can be approximated by

**Figure C.12:**
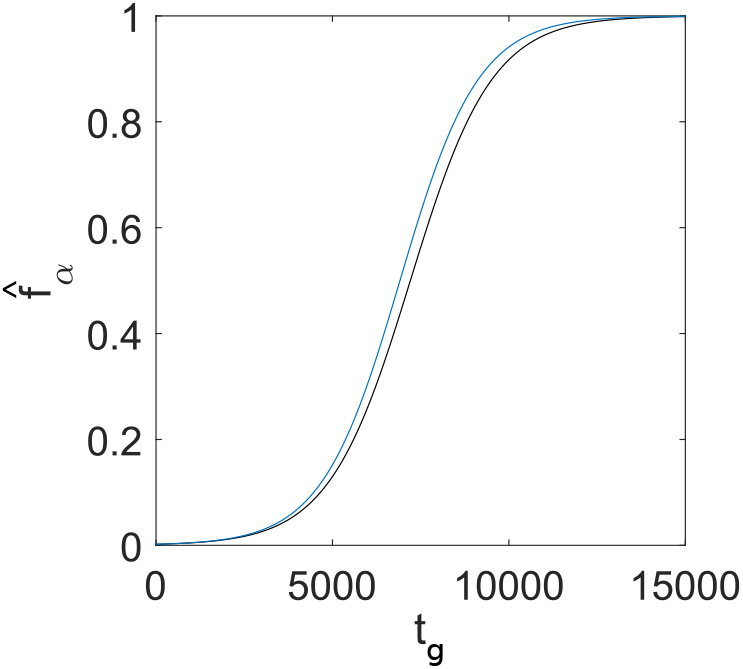
Invasion dynamics for a mutant with fertilization rate *α* + *δα*. Blue - analytical prediction using Eq. (C.2), black - numerical simulation. The initial conditions and parameters are the same as in Figure C.11 except *δα* = 1/200, *δm* = 0 and *G* = 1.5 × 10^4^.

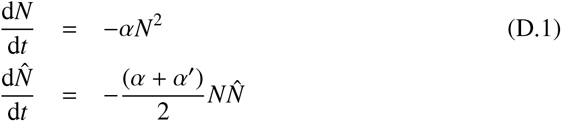

with *N*(0) = (*AM*)/*m* and 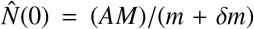 if a mutation occurs in *m*. As mutants are assumed to be rare, we have here neglected mutant-mutant interactions which significantly simplifies the analysis [34, 93] (see Eq. (B.5) for comparison). We now consider the fitness of the mutant, where the mutant has mass 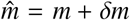 and fertilisation rate 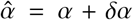. The fitness of the mutant relative to that of the resident is given by

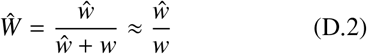

where *w* and 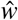 are the absolute fitness of the mutant and resident respectively and the approximation for the relative fitness 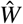 holds to leading order when mutants are rare. The absolute fitnesses can be derived from Eq. (D.1) as

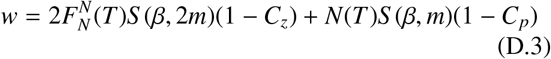

and

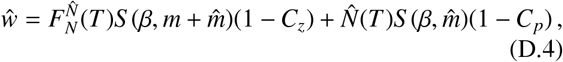

with 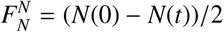 and 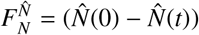 To derive the evolutionary ODEs, we calculate the fitness gradient given by

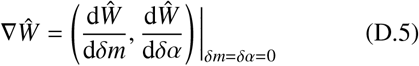

which gives exactly what is given in Eq. (5). Two specific scenarios of Eq. (D.2) can be simply obtained by linearising Eq. (B.11) or Eq. (B.13) about their mutant free steady states 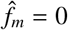 or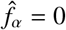, i.e

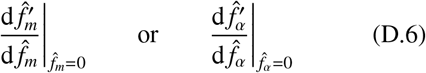

Since Eq. (B.11) and Eq. (B.13) considers mutations in one trait at a time, 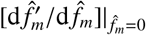 in Eq. (D.6) equals Eq. (D.2) evaluated at *δα* = 0 and 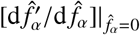 equals Eq. (D.2) evaluated at *δm* = 0. We prove this as follows. Denoting 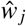 as the absolute fitness where 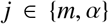, Eq. (B.11) or Eq. (B.13) can be expressed as

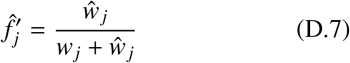

and applying the quotient rule, we find

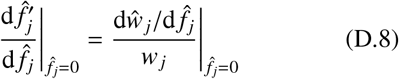

since 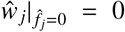 By noting that 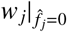 is equal to Eq. (D.3) and 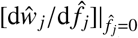 is equal to Eq. (D.4) with *δm* or *δα* equal to 0. This shows how this multidimensional approach yields the exact same evolutionary ODEs, Eq. (5), as is given in the main text.

The following appendix gives an analogous derivation using instead the invasion dynamics that assume mutations in *m* and *α* occur independently, however this does not change the final evolutionary dynamics obtained.

## Appendix E. Evolutionary dynamics: Fixed environment

### Appendix E.1. Derivation of evolutionary ODEs

We begin by noting that the functional form of 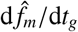 (see Eq. (C.1)) and 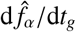 (see Eq. (C.2)) is 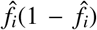, which implies a situation of trait substitution [67]; mutants are either driven to fixation or extinction, and polymorphic equilibria are not possible. This simplifies the subsequent analysis considerably.

Taking a classic adaptive dynamics approach [67] and define the invasion fitness of the mutants as their percapita rate of reproduction upon arising in the population (i.e. when 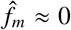 and 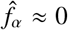). Under the standard assumptions of adaptive dynamics (i.e. that mutations are of small effect, 1≫ *δm, δα*, and occur sufficiently rarely that each mutation can fixate before a new mutation occurs), the evolutionary dynamics are given by

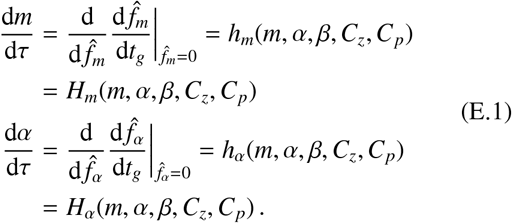

Substituting for 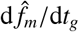 from Eq. (C.1), and 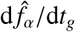 from Eq. (C.2), we obtain Eq. (5) in the main text.

The evolutionary dynamics given in Eq. (E.1) can also be derived using the multidimensional approach as described in [91, 90] (see Appendix D).

### Appendix E.2. Analysis of ODEs

In this section we aim to analytically characterise the long term evolutionary behaviour of the population in a fixed environment, with dynamics given by Eq. (5), as illustrated in Figure 3.

We begin by calculating the evolutionary behaviour of *m* when the fertilization rate is fixed to zero (*α* = 0). Solving *H*_*m*_(*m*, 0, *β, C*_*z*_, *C*_*p*_) = 0, for 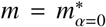 we obtain

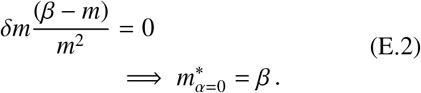

We now turn to the evolutionary behaviour of *m* and *α*. First, we substitute the point (*m, α*) = (*β*, 0) calculated in Eq. (E.2) into Eq. (E.1), which gives

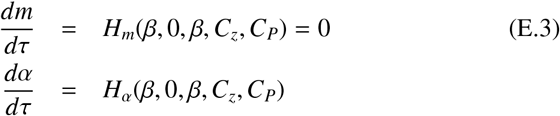

For 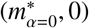 identified above to remain stable if evolution on *α* is allowed requires that [d*α*/d*τ*] | _(*m*=*β,α*=0)_ < 0 (i.e. that evolution selects against increases in *α*). Stating this condition in full, we have

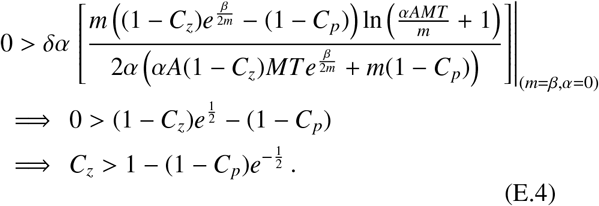

If fertilization costs exceed this value, evolutionary trajectories starting at 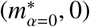 will remain there (see Figure 3, panel (b)). This can be shown analytically by checking that it is an evolutionary stable state. First, we prove that it is convergence stable by showing that *∂*_*m*_[*dm*/*dτ*]|_(*m*=*β,α*=0)_ < 0.

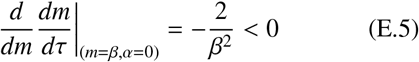

which implies that (*m, α*) = (*β*, 0) is stable. The evolutionary stability of this point will be proven in Appendix H. Conversely if fertilization costs do not exceed this value, evolutionary trajectories starting at 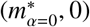 will initially experience a selective pressure for increasing *α*, due to the fact [d*α*/d*τ*] | _*m*=*β*_ > 0 under these conditions (see Figure 3, panel (a)); the dynamics are then pulled to another evolutionary singularity, which we characterise below.

When *C*_*z*_ > 1− (1−*C*_*p*_) exp (−1/2) (but less than another critical value, yet to be determined), only a subset of initial conditions fall within the basin of attraction of this fixed point described above (see Figure 3, panel (b)), with remaining initial conditions leading to an evolutionarily state at which *α*^∗^ → ∞. Conversely, when *C*_*z*_ < 1− (1− *C*_*p*_) exp (− 1/2), *all* initial conditions lead to this fixed point at which *α*^∗^→ ∞ (see Figure 3, panel a). We now calculate the mass to which the population evolves at this second fixed point. Taking the limit *α* → ∞ in the evolutionary dynamics for *m* (see Eq. (5)) and solving for zero;

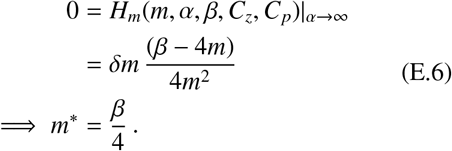

Therefore the second early fixed point is at (*m, α*) → (*β*/4, ∞).

Since there is likely an upper limit on the fertilisation rate biologically *α*_max_, an alternative approach is to conduct a stability analysis about (*m, α*) = (*β*/4, *α*_max_). Furthermore, due to the small selection strength to increase *α* when *α* is sufficiently high, *α*_max_ can be thought of as a quasi-equilibrium. For the parameters in Figure 3, the selection strength to increase *α* |*H*_*α*_(*m, α, β, C*_*z*_, *C*_*p*_) | becomes 6.86×10^−5^ when *α* = 10. To determine the stability of Eq. (5), we calculate the eigenvalues of its Jacobian evaluated at (*m, α*) = (*β*/4, *α*_max_). The functional form of all entries of the Jacobian is too lengthy to include in the manuscript, however it is included in the *Mathematica* code provided in the Data availability statement. For the parameter values in Figure 3, the eigenvalues of the Jacobian evaluated about (*m, α*) = (*β*/4, 10) are −16.00 and −1.29 × 10^−5^, which implies it is convergence stable.

For even greater costs of fertilization, *C*_*z*_, our mathematical analysis (which assumes monomorphic resident populations and no evolutionary branching) suggests that 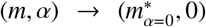 becomes the only attractor, with (*m, α*) → (*β*/4, ∞) ceasing to be a fixed point. To determine the critical cost at which this occurs, we take [d*α*/d*τ*]_*m*=*β*/4_ and calculate the conditions under which this is negative when *α* is large (i.e. when (*m, α*) → (*β*/4, ∞) is no longer attracting, but repelling). Expanding [d*α*/d*τ*]_*m*=*β*/4_ in small 1/*α*, we find that to leading order we must have;

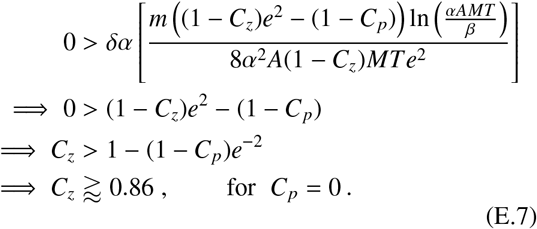

However, while this accurately captures the short-term evolutionary dynamics, we see evolutionary branching (which our model Eq. (5) does not account for) should trajectories approach the *m* ≈ *β*/4 manifold. Unless costs are exceedingly high (*C*_*z*_ ≈ 1), this eventually leads to anisogamy followed by oogamy (see Figure E.13). We discuss this more in Appendix H.

### Appendix E.3. Implementation of simulations

Here we detail the process of numerical simulation (see *Code availability statement*). We employ a multigenotype model whereby a mutation occurs at rate *µ*. This rate is the inverse of the expected number of fertilization processes (i.e. generations) until the next mutation event. Before each successive mutation event, the number of fertilization processes until the next mutation event is determined by generating a number from a *Geo*(*µ*) distribution and taking the inverse of that number. In an *S* genotype model, if a given genotype *i* has a trait value of (*m*_*i*_, *α*_*i*_) and has frequency *f*_*i*_, then the mean population trait value is given by

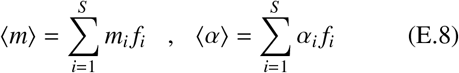

The simulations in Figure 3 are repeated for 5500 mutation events. Below, we detail how we simulate the dynamics on each timescale.

#### Fertilization Kinetics

We construct our model for fertilization kinetics assuming that each genotype, characterised by their unique (*m*_*i*_, *α*_*i*_), can fertilize with one another. The input parameters of this function are the trait values of each genotype ***m*** and *α*, their frequency in the preceding adult generation ***f***, and the parameters *A, M* and *T*. The fertilization kinetics simulations are then run according to

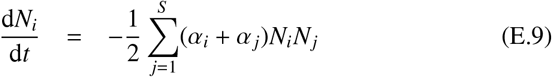

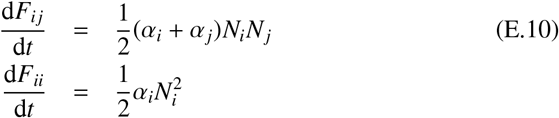

where *F*_*i j*_ is the number of fertilized cells formed from genotypes *i* and *j*, and *F*_*ii*_ is the number of fertilized cells formed from two cells of genotype *i* for any *i, j* ∈ [1, *S* ]. The initial conditions are

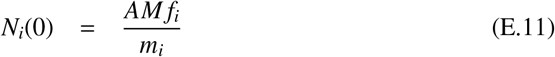

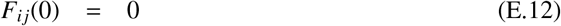

The fertilization process is run for a fixed time period *T*. At the end of this time period, the function outputs the number of cells of each type. These include the number of unfertilized cells with each trait pair, *N*_*i*_(*T*), and fertilized cells of each type *F*_*i j*_(*T*).

#### Single Generation Dynamics

We simulate the frequency of each genotype after the end of each generation taking into account the survival probability of each genotype. Upon maturation, only a fraction of unfertilized and fertilized cells survive to adulthood. We use the outputs of the fertilization kinetic function along with the Vance survival functions Eq. (2) to calculate the probability that each progeny survives into adulthood. The single generation dynamics are run according to

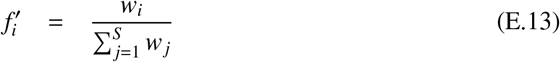

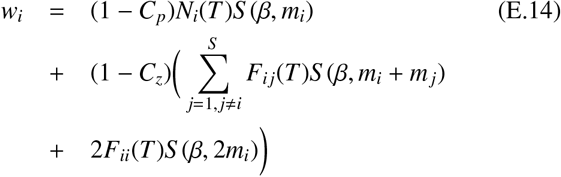

where 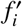 is the frequency of genotype *i* in the subsequent generation and *w*_*i*_ is its absolute fitness. The input parameters of this function are *N*_*i*_(*T*) and *F*_*i j*_(*T*), ***m***, *α, β, C*_*z*_ and *C*_*p*_ whilst the output is ***f***, the frequency of all genotypes at the end of the generation.

#### Invasion Dynamics

The invasion dynamics are run for approximately one unit of *τ* (i.e. until the next randomly chosen mutation event after *G* generations). As outlined in the opening paragraph of Appendix E.3, *G* is generated from a *Geo*(*µ*) distribution following each mutation event. The inputs of this function are ***f***, ***m***, *α, A, M, T, C*_*z*_, *C*_*p*_, *G* and *β* and the output is the frequency of each genotype ***f*** after one unit of *τ* (i.e after *G* fertilization processes). Please note we have adopted one single notation for the frequency of all genotypes.

#### Evolutionary Dynamics

The evolutionary dynamics have the input parameters *δ, µ*, ***f***, Nmut, *f*_0_, *A, M, T, C*_*z*_ and *C*_*p*_ where *δ* is the mutational stepsize, the initial frequency of a newly introduced mutant *f*_0_ is chosen to be small (equal to 0.002 in our model). We initialise the simulation with two genotypes, where each genotype is characterised by a unique pair of trait values e.g. (*m*_*i*_, *α*_*i*_) for genotype *i*. A mutant is introduced into the population after a random number of fertilization processes *G* (generated from a *Geo*(*µ*) distribution) at frequency *f*_0_. The mutation is chosen to occur in either *α* or *m* with equal probability 1/2. The mutation also acts to increase/decrease the trait value each with probability 1/2. Upon introduction of this mutant, the invasion dynamics of the population is run for *G* fertilization processes. In the meantime, the mean mass and fertilization rate of the population is recorded using Eq. (E.8). Next, we repeat the process of introducing a new mutant into the population. Since the population now has more than two genotypes, the genotype that mutates is chosen with probability weighted by the frequency of each genotype. There is now the possibility of back mutation to one of the existing genotypes. In this case, if an existing genotype *k* with frequency *f*_*k*_ mutates to another existing genotype *l* that has frequency *f*_*l*_, then following mutation, the frequency of genotype *k* becomes *f*_*k*_− *f*_0_ and the frequency of genotype *l* becomes *f*_*l*_ + *f*_0_. Furthermore, a genotype is thought to be extinct if its frequency falls below 10^−3^, in which case we remove that genotype.

#### Appendix E.4. Evolutionary Branching in Gamete Mass and fertilization rate

In this section we present additional numerical results investigating the evolutionary branching that occurs in simulations on the manifold *m* ≈ *β*/4 (along which d*α*/d*t* ≈ 0).

##### In a fixed environment, when C_z_ is small relative to C_p_, branching in fertilization rate still occurs but no longer gives rise to oogamy

When the cost to fertilization, *C*_*z*_ is small compared to *C*_*p*_, we find that although branching in both gamete mass and fertilization rate still occurs, the branching in *α* no longer acts to decrease the fertilization rate of macrogametes. Therefore, we observe pseudooogamy but not oogamy. This behaviour is illustrated in Figures 6 and I.19. We can estimate the critical value of *C*_*z*_ at which this occurs by comparing the survival probability of a macrogamete that does not fertilize with a microgamete, which has cost *C*_*p*_ with that of a macrogamete that does fertilize with a microgamete at cost *C*_*z*_; if this first probability exceeds the second, there should be no evolutionary pressure for oogamy to evolve. We find

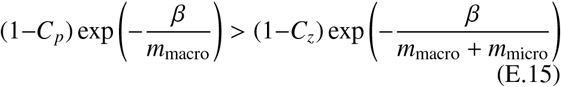

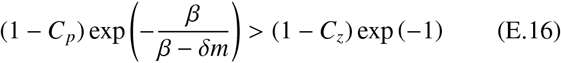

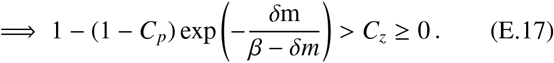

##### In a fixed environment, when C_z_ is very large, branching in mass and fertilization rates can still occur, with oogamy possible (dependent on initial conditions)

Our analysis of the dynamics of Eq. (5) in Appendix E.2 suggested that zero fertilization rates is the only evolutionary attractor when costs to fertilization are high (see Eq. (E.7)). However, our analysis in Appendix H along with our simulations reveal that although d*α*/d*t* < 0 along the *m*≈ *β*/4 manifold, any trajectory that approaches this manifold can experience branching in gamete mass, unless *C*_*z*_ = 1. Once branching in gamete mass occurs, the smaller gametes (e.g. microgametes) again experience a strong selective pressure to increase their fertilization rate, despite the high costs imposed by fertilization; sexual conflict drives the population towards obligate oogamy imposed by motile microgametes. Thus as for *C*_*z*_ < 1, there is always a subset of initial conditions that lead to trajectories along the *m* ≈ *β*/4 manifold (most obviously initial conditions on the manifold itself) and thus oogamy remains one of the two evolutionary outcomes, albeit requiring an increasingly small and biologically unrealistic set of initial conditions (see Figure E.13 below).

**Figure E.13:**
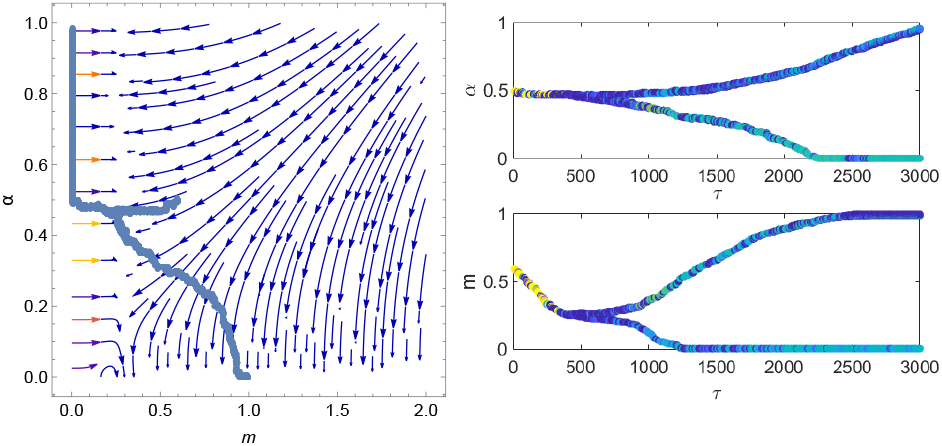
Numerical illustration of evolutionary branching for the case where *C*_*z*_ = 0.9 and *C*_*p*_ = 0. All other parameters the same as Figure 3, except (*m*(0), *α*(0)) = (0.6, 0.5) and run for 3000/µ generations.

## Appendix F. Evolutionary dynamics: switching environments with bet-hedging

We first tackle the derivation of the approximate dynamics in the FRTI (fast relative to invasion) switching regime in Appendix F.1, before verifying the qualitative robustness of these results in the FRTE switching regime in Appendix F.1.

### Appendix F.1. Derivation of evolutionary ODEs: FRTI

When the environment switches between the two environments many times before an invasion has time to complete, the selection pressure would alternate between that of the two environments rapidly and the population effectively experiences the weighted average of the two environments. The invasion dynamics can be approximated by multiplying the frequency dynamics by the weighted average of the selection gradients of the two environments;

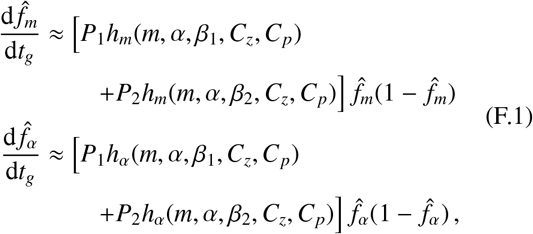

where *h*_*m*_(*m, α, β, C*_*z*_, *C*_*p*_) and *h*_*α*_(*m, α, β, C*_*z*_, *C*_*p*_) are given in Eq. (C.1) and Eq. (C.2) respectively. In Figure F.14 we show that this indeed is a good approximation of the dynamics when *δm* and *δα* are small and when the residency times in each environment are small relative to the invasion time. Note that as in the case of the fixed environment, the functional form of *f*_*i*_ in these equations implies that trait substitution occurs for independent mutations on the gamete mass and fertilization rate.

**Figure F.14:**
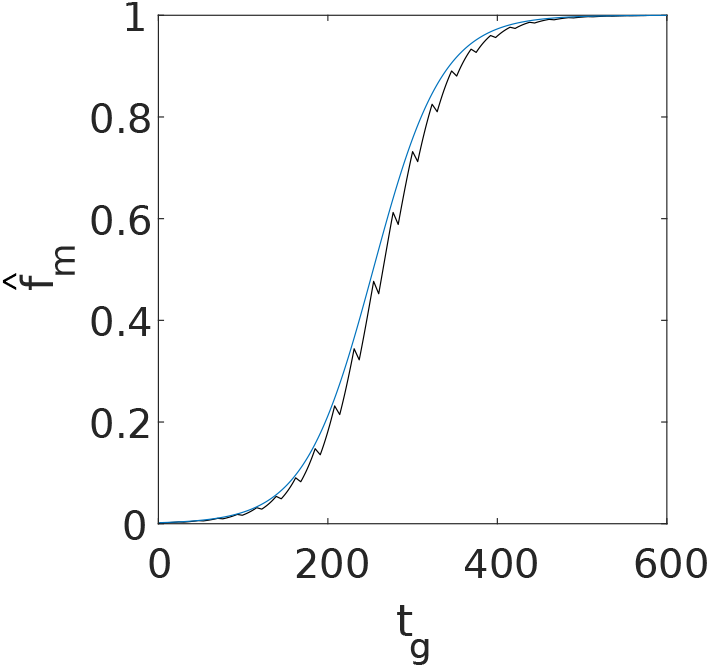
Invasion dynamics for a population undergoing bet-hedging when the environment switches FRTI. Blue curve is the analytical approximation using Eq. (F.1) and jagged curve is the numerical simulation. Mutation occurred in mass with *δ* = 0.005, *f*_0_ = 0.002, (*m*(0), *α*(0)) = (0.3, 0.1), λ_1→2_ = 13/222, λ_2→1_ = 1/6. All other parameters the same as Figure 7.

We now apply the same approach as in Appendix E.1 (see Eq. (E.1)) to obtain

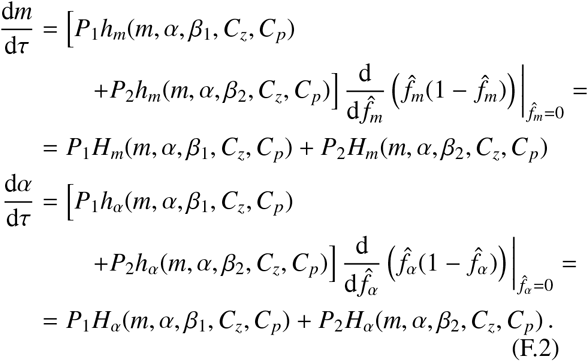

We see in Figure 7 that these also provide a good approximation of the evolutionary dynamics.

### Appendix F.2. Derivation of evolutionary ODEs: FRTE

In the FRTI scenario in the previous section, we supposed that switching between the environments was happening sufficiently regularly relative to the timescale of invasion that the effective invasion dynamics could be described by a weighted mean of the invasion dynamics in both environments (see Eq. (F.1)). Applying an analogous logic, we now assume in the FRTE scenario that switching between the environments occurs regularly relative to the timescale of evolution (the arrival rate of new mutations) that the effective evolutionary dynamics can be described by a weighted mean of the evolutionary dynamics in both environments; that is

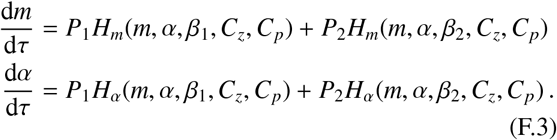

We note that these are in fact exactly the same evolutionary dynamics as derived in the FRTI scenario (see Eq. (F.2)). In Figure 7 we show that these do indeed remain a good approximation for the dynamics in the FRTE scenario. The difference between the dynamics in both regimes is quantitative, rather than qualitative. In the very-fast switching FRTI regime, the populations follow the effective dynamics very closely. In the comparatively slower FRTE regime, although the populations no longer follow the dynamics as well, the qualitative picture of the dynamics is still captured by Eq. (F.3). In particular, we still observe an evolutionarily stable state of intermediate fertilization rate (see Figure 7).

### Appendix F.3. Analysis of ODEs

We begin, as in Appendix E.2, by considering the evolutionary behaviour of *m* when the fertilization rate is fixed to zero (*α* = 0). We recall that if the population was fixed in environment 1 or 2 with *α* = 0, the mass of gametes would evolve to *m* = *β*_1_ and *m* = *β*_2_ respectively. In the switching environment we instead find the bet-hedging strategy

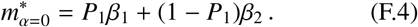

**Figure F.15:**
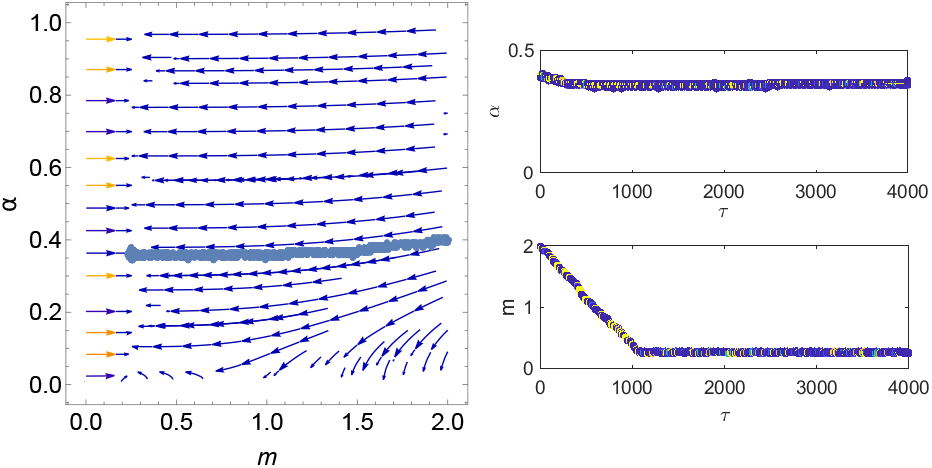
Numerical simulation showing an absence of branching for a system undergoing bet-hedging in an environment that switches FRTI. Parameters are same as Figure 7 (a) and system run for 4000/µ generations.

Meanwhile the region of the boundary *α* = 0 over which reduced fertilization rates are selected for is given by [d*α*/d*τ*]|_*α*=0_ < 0, or

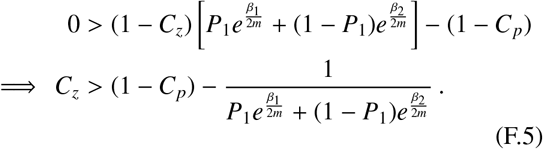

If this condition holds 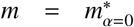, then the switching-induced fixed point 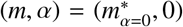 is stable. We next turn to the high fertilization rate fixed point, for which *α* → ∞. In a similar manner to Eq. (E.6) (albeit with *H*_*m*_(*m, α, β, C*_*z*_, *C*_*p*_) replaced with [*P*_1_*H*_*m*_(*m, α, β*_1_, *C*_*z*_, *C*_*p*_) + *P*_2_*H*_*m*_(*m, α, β*_2_, *C*_*z*_, *C*_*p*_)]), we find the bet-hedging strategy

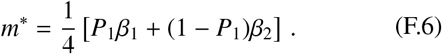

However, unlike in the fixed environment case, (*m, α*) = (*m*^∗^, ∞) is not always a fixed point.

In the switching environment, a third evolutionary attractor can emerge, brought about by a balance between selection for high fertilization rates in one environment and zero fertilization rates in the other. Solving *P*_1_*H*_*α*_(*m, α, β*_1_, *C*_*z*_, *C*_*p*_) + *P*_2_*H*_*α*_(*m, α, β*_2_, *C*_*z*_, *C*_*p*_) = 0 for *α* we obtain

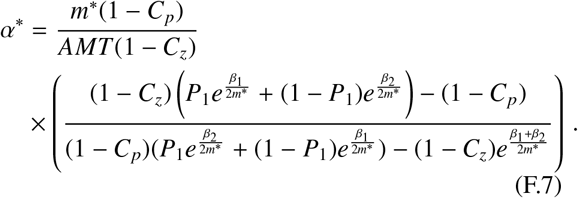

A good approximation for *m*^∗^ in this equation can be deduced by noting that the fixed point sits on a vertical manifold of trajectories along which *m* is approximately held constant as *α* → ∞; thus we substitute ≈ *m*^∗^ from Eq. (F.6) to obtain the approximate expression for the attractor (*m*^∗^, *α*^∗^) given in Eq. (16). In Appendix F.5, we verify from simulations that evolutionary branching does not occur at this switching-induced fixed point, and that the population is instead held in a state of isogamy.

### Appendix F.4. Implementation of simulations

In the bet-hedging scenario, we simulate the evolutionary dynamics by implementing a Gillespie algorithm. In particular, we introduce mutations and environmental switching events randomly with geometrically distributed waiting times, where the waiting time is measured in units of number of fertilization processes *t*_*g*_. To simulate the FRTI regime, we set the environmental switching rates to larger values than the mutation rate i.e. *λ*_2→1_, *λ*_1→2_ >> *µ*. Likewise, in the FRTE regime, we set *λ*_2→1_ and *λ*_1→2_ to smaller values than *µ*. The mutation rate in the numerical simulation of the FRTI regime overlaid in Figure 7 (a) is *µ* = 3.5 × 10^−4^ and the switching rates are *λ*_2→1_ = 67/532 and *λ*_1→2_ = 1/4. For Figure 7 (b) we have *µ* = 3.5 × 10^−4^ and the switching rates are *λ*_2→1_ = 1/6 and *λ*_1→2_ = 13/222. For the FRTE regime, *µ* = 5 × 10^−4^ and the switching rates are *λ*_2→1_ = (67*µ*)/532 and *λ*_1→2_ = *µ*/4 in Figure 7 (a) and *λ*_2→1_ = *µ*/6 and *λ*_1→2_ = (13*µ*)/222 in Figure 7 (b).

**Figure F.16:**
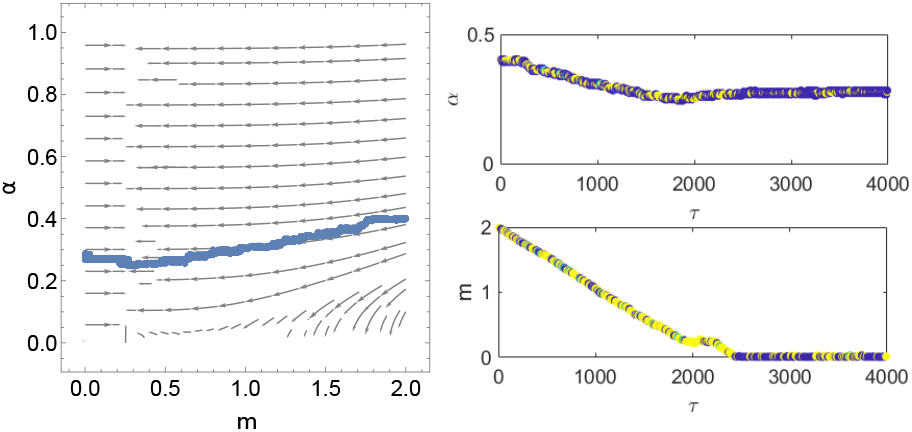
Numerical illustration showing an absence of evolutionary branching for a system undergoing bet-hedging in an environment that switches FRTE. Parameters the same as Figure 7 (c) and the system is run for 4000/µ generations.

### Appendix F.5. Absence of Evolutionary Branching at Switching-Induced fixed point

In this section we present additional numerical results that confirm that evolutionary branching does not oc-. cur at the switching-induced fixed point calculated in Eq. (16). We see that under both FRTI and FRTE, the population is held in a state of isogamy. This behaviour is illustrated in Figures F.15 and F.16.

### Appendix F.6. Interpretation of Mutant Frequency Growth Rate in Switching Environments

Eqs. (C.1) and (C.2) give the invasion dynamics in a fixed environment. For simplicity, suppose we consider the dynamics when the mutant frequencies, 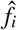, are small, as is the case at the beginning of an invasion. The mutant frequencies then increase exponentially such that

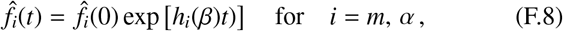

where we have suppressed the dependence of *h*_*i*_ on other parameters for clarity.

When environmental switching is taking place according to a telegraph process, the time spent in each environment becomes a random variable. Suppose we consider *S* switching events, beginning in environment 1. The times spent in environment 1 is *T*_1, *j*_ ∼ exp(*λ*_1→2_) during the *j*^*t*^*h* period in environment 1, and similarly for environment 2 (*T*_2, *j*_ ∼ exp(*λ*_2→1_)). The increase in mutant frequency is then given by

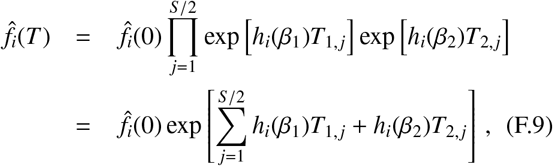

where 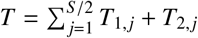 The sums over *T*_1, *j*_ and *T* _2, *j*_ in the above equations result in two Gamma-distributed random variables in the exponent. Calculating the expected value of Eq. (F.9) is thus not trivial [72]. However, we can gain some intuition by expressing the exponent in Eq. (F.9) in terms of an expansion about the mean:

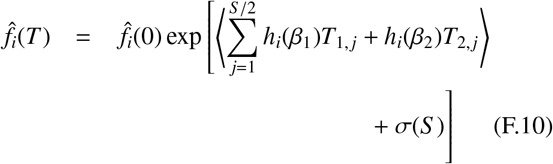

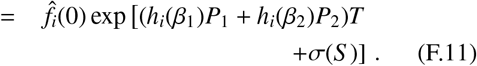

where *σ*(*S*) is the standard deviation of the exponent in Eq. (F.9) and (*h*_*i*_(*β*_1_)*P*_1_ + *h*_*i*_(*β*_2_)*P*_2_)*T* is the expectation (see Eq. (7)). We expect the standard deviation *σ*(*S*) to be a decreasing function of the number of switching events *S* (or equivalently of the switching rates *λ*_1→2_) and *λ*_2→1_) if one considers some fixed time *T*) by the law of large numbers. Therefore fast switching reduces *σ*(*S*), and thus the role variance in growth rates between environments [73, 72] is muted in this model setup. While this argument is borne out by the success of the approximation in predicting the population dynamics (as illustrated in Figs F.14-F.16), it would be interesting to explore this more formally mathematically as the generalisation of geometric mean fitness is often model-specific [94].

## Appendix G. Effect of minimum microgamete mass on macrogamete mass

Here, we provide a simulation of what happens to the macrogamete mass if we impose a minimum mass on the microgamete.

In Figure G.17, we see that as the macrogamete mass *m*_*macro*_ becomes sufficiently large, the macrogamete evolves to decrease *α*, avoiding fertilisation costs. As *α* approaches near 0, *m*_*macro*_ begins to increase sharply, tending towards *m*_*macro*_ ≈*β*− *m*_*micro*_. This suggests that on the *α* = 0 boundary, the zygote is expected to evolve a mass of *β*.

**Figure G.17:**
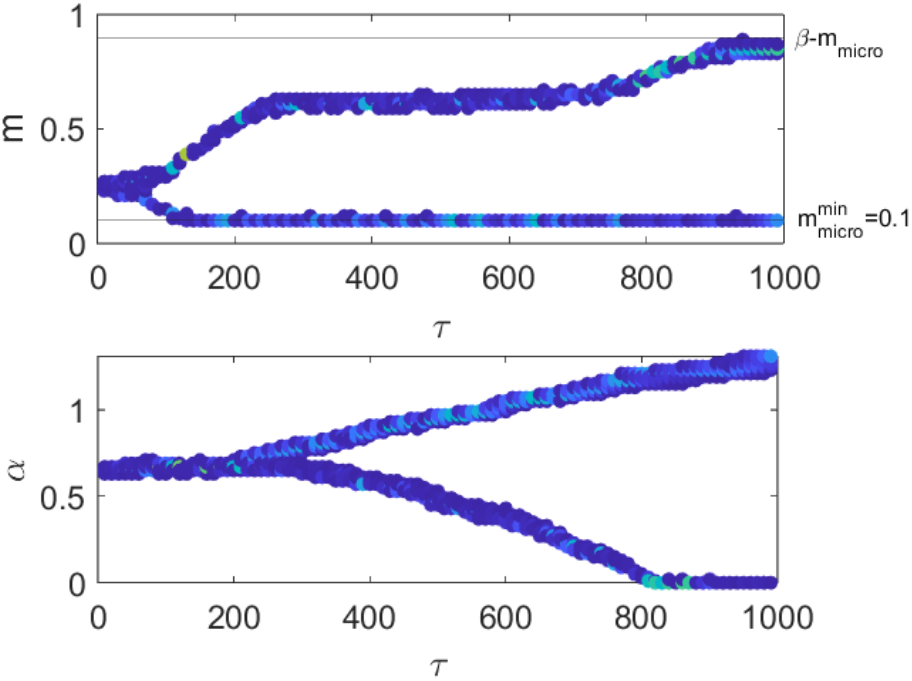
Evolutionary branching simulation when we impose a minimum microgamete size 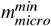 larger than *δm* (Top - evolution of *m*, bottom - evolution of α). Parameters are as in Appendix J but 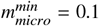 and (*m*(0), *α*(0)) = (0.25, 0.7).

## Appendix H. *α*_max_ below which branching to anisogamy can be arrested

The two necessary conditions for evolutionary branching to occur is for the fitness gradient of an invading mutant to be zero (i.e. *dm*/*dτ* in Eq. (E.1) to equal zero if the mutation occurs in mass, this is also known as an evolutionary singularity) and the second derivative of Eq. (B.11) with respect to the mutational stepsize *δm* to be positive [95, 96] when *δm* = 0. Using Eq. (E.1), we see that the vertical manifold *m*≈ *β*/4 is an approximate evolutionary singularity since [*dm*/*dτ*] |_*m*=*β*/4_≈ 0. The condition for selection in mass to be disruptive about an evolutionary singularity is

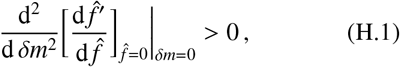

where 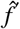 is the frequency of the mutant in the subsequent generation, given by Eq. (B.11). The explicit expression for the left hand side of Eq. (H.1) is given by

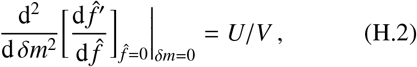

where

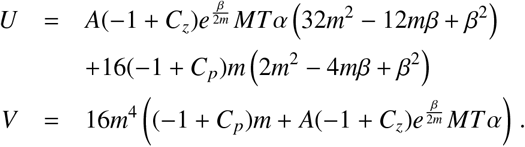

The first condition for branching to occur in mass i.e *dm*/*dτ* = 0 can be calculated straightforwardly by setting Eq. (E.1) to zero and solving for *α*, which gives

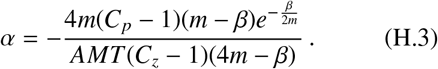

We then substitute Eq. (H.3) into Eq. (H.2) to obtain

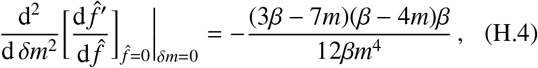

which needs to be positive for selection to be disruptive. Eq. (H.4) equals 0 if *m* = 3*β*/7 or *m* = *β*/4 and is positive for *β*/4 < *m* < 3*β*/7. By substituting the boundaries of this interval into Eq. (H.3), we find that it corresponds to

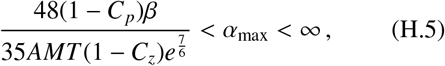

and thus the value of *α*_max_ above which branching will occur in mass is the value given on the left hand side of the inequality Eq. (H.5). In other words, Eq. (H.5) is the interval in *α*_max_ in which branching is expected to occur. The vertical line in Figure 5 corresponds to the lower limit of this interval. In Figure H.18 below, we provide a numerical example showing how isogamy can be stabilized below a sufficiently low *α*_*max*_.

**Figure H.18:**
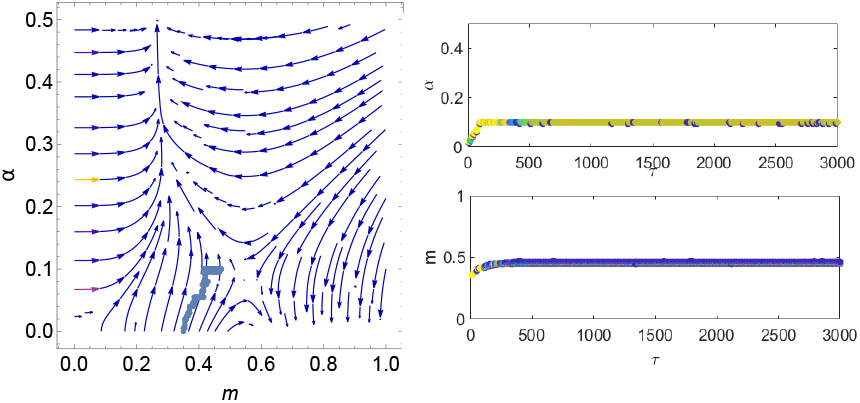
Numerical illustration of the stabilization of isogamy below a sufficiently low α_*max*_. System parameters are *A* = 100, *M* = 1, *T* = 0.1, *C*_*z*_ = 0.6, *C*_*p*_ = 0, *β* = 1, *α*_*max*_ = 0.1 and simulation parameters are *δ* = 5 ×10^−3^, *f*_0_ = 2 ×10^−3^ and run for 6 ×10^6^ generations. Using Eq. (H.5) we can calculate that branching would occur if *α*_*max*_ ⪆ 0.1068 for these parameter values.

## Appendix I. The stabilization of Anisogamy under high costs of parthenogenesis relative to fertilization

In Figure I.19, we provide an example of a parameter regime where pseudooogamy occurs. When the parameters are close to the boundary where we observe the transition between oogamy and anisogamy (i.e close to where the inequality in Eq. (E.17) becomes an equality), we observe pseudooogamy, where there is a considerably stronger selection pressure for microgametes to increase their fertilization rate than macrogametes. Under pseudooogamy, macrogametes are still motile but are considerably less motile than microgametes.

**Figure I.19:**
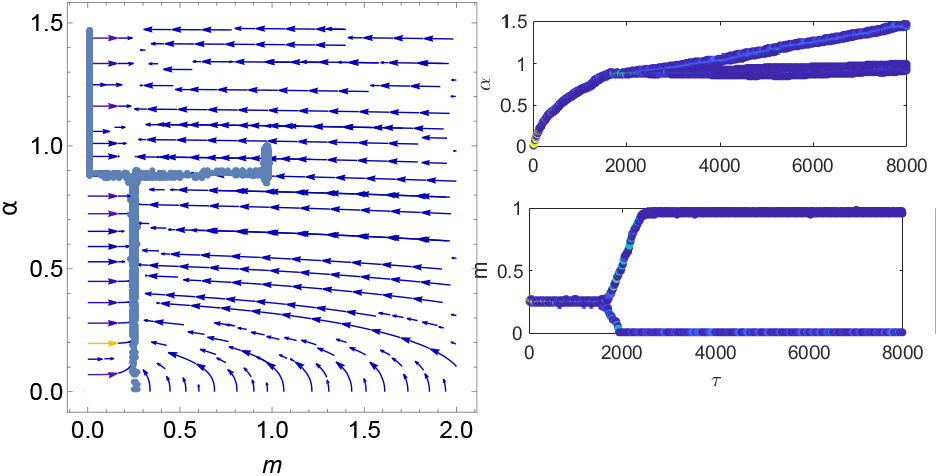
Numerical illustration of the evolution of pseudo-oogamy in fixed environment. Parameters are *C*_*z*_ = 0, *C*_*p*_ = 0, *δm* = 0.005, (*m*(0), *α*(0)) = (0.25, 0) and simulation run for 1.6× 10^7^ generations. Remaining parameters are as given in Appendix J.

## Appendix J. Model parameters in figures

### Appendix J.1. Figure 3

The initial points of the trajectories are (*m*(0), *α*(0)) = (1.5, 0.6) and (*m*(0), *α*(0)) = (2, 0.1). Remaining model parameters are *A* = 100, *M* = 1, *T* = 1 and *β* = 1. Simulation parameters: Initial frequency of novel mutant genotype *f*_0_ = 2 × 10^−3^, mutation rate *µ* = 5 × 10^−4^ (number of generations)^−1^, run for 1.1 × 10^7^ generations in panel (a) and 1.24 × 10^7^ generations in panel (b). The mutational stepsize is *δ* = 5 × 10^−3^. Throughout the paper, we use *δm* and *δα* to denote mutational stepsize in *m* and *α* respectively, however in simulations involving co-evolution of *m* and *α*, the same stepsize is used for both traits. For convenience, we thus denote mutational stepsize as *δ* in these simulations.

## CRediT authorship contribution statement

**Xiaoyuan Liu:** Methodology, Formal analysis, Investigation, Software, Writing- original draft preparation. **Jonathan W. Pitchford:** Conceptualization, Supervision, Writing-Reviewing and Editing. **George W. A. Constable:** Conceptualization, Methodology, Supervision, Formal analysis, Writing - review & editing.

## Declaration of interest

Declaration of interest: none.

## Acknowledgements

This work was supported in part by the Ph.D. studentship funded to Xiaoyuan Liu by the University of York though the EPSRC DTP, Grant No. EP/V52010X/1.

## Data availability

This manuscript did not use any real data. All the codes required for generating the simulation data in the manuscript are provided at https://github.com/phil-liu1020/ADSW origin sexes, and is described in Appendix E.3 and Appendix F.4

## Notes

https://github.com/phil-liu1020/ADSW_origin_sexes

### Competing Interest Statement

The authors have declared no competing interest.

### Summary of Updates

This manuscript has undergone a round of major revision in response to a peer review. This has been resubmitted lately and we hereby upload the revised manuscript.

